# Myeloid cell interferon responses correlate with clearance of SARS-CoV-2

**DOI:** 10.1101/2021.06.28.450153

**Authors:** Dhiraj K. Singh, Ekaterina Aladyeva, Shibali Das, Bindu Singh, Ekaterina Esaulova, Amanda Swain, Mushtaq Ahmed, Journey Cole, Chivonne Moodley, Smriti Mehra, Larry S. Schlesinger, Maxim N. Artyomov, Shabaana A. Khader, Deepak Kaushal

## Abstract

The emergence of mutant SARS-CoV-2 strains associated with an increased risk of COVID-19-related death necessitates better understanding of the early viral dynamics, host responses and immunopathology. While studies have reported immune profiling using single cell RNA sequencing in terminal human COVID-19 patients, performing longitudinal immune cell dynamics in humans is challenging. Macaques are a suitable model of SARS-CoV-2 infection. We performed longitudinal single-cell RNA sequencing of bronchoalveolar lavage (BAL) cell suspensions from adult rhesus macaques infected with SARS-CoV-2 (n=6) to delineate the early dynamics of immune cells changes. The bronchoalveolar compartment exhibited dynamic changes in transcriptional landscape 3 days post-SARS-CoV-2-infection (3dpi) (peak viremia), relative to 14-17dpi (recovery phase) and pre-infection (baseline). We observed the accumulation of distinct populations of both macrophages and T-lymphocytes expressing strong interferon-driven inflammatory gene signature at 3dpi. Type I IFN response was highly induced in the plasmacytoid dendritic cells. The presence of a distinct HLADR^+^CD68^+^CD163^+^SIGLEC1^+^ macrophage population exhibiting higher angiotensin converting enzyme 2 (ACE2) expression was also observed. These macrophages were significantly recruited to the lungs of macaques at 3dpi and harbored SARS-CoV-2, while expressing a strong interferon-driven innate anti-viral gene signature. The accumulation of these responses correlated with decline in viremia and recovery. The recruitment of a myeloid cell-mediated Type I IFN response is associated with the rapid clearance of SARS-CoV-2 infection in macaques.

## Introduction

The underlying immune mechanisms that drive disease versus protection during the Coronavirus disease 2019 (COVID-19) are not well understood. Analysis of system-wide transcriptomic responses can be extremely useful in identifying features of protection and pinpoint the host immune processes involved in the control of infection and drivers of pathology^1,2^. Transcriptional changes in cells in the broncho-alveolar lavage (BAL) and peripheral blood mononuclear cells (PBMCs) of COVID-19 patients show distinct host inflammatory cytokine profiles, suggesting that excessive cytokine release is associated with COVID-19 pathogenesis ^3^. However, analyses were conducted using end-point samples in patients, and it is possible that the excessive cytokine storm at that time is a representation of an exacerbated viral infection and associated immune dysregulation. We recently developed a nonhuman primate (NHP) model of SARS-CoV-2 infection^4^, where NHPs develop signs of COVID-19 disease including characteristic ground glass opacities in lungs, coinciding with a cytokine storm and a myeloid cell influx, followed by clearance of the virus and recovery^4^. Using RNAseq, we showed that Interferon (IFN) signaling, neutrophil degranulation and innate immune pathways were significantly induced in the SARS-CoV-2-infected lungs of NHPs, while pathways associated with collagen formation were downregulated ^5^. Since these animals controlled infection naturally, our results point to the importance of early innate immune responses and cytokine signaling, particularly Type I IFN signaling, in protecting against COVID-19. One limitation of the above study was that it was conducted in terminal lung samples and thus may not represent the dynamic changes that occur immediately after infection. Furthermore, system-wide transcriptomics was studied using bulk-RNAseq, thus averaging the overall contributions of various cell types and pathways at play.

The use of single-cell technologies such as RNA-sequencing (scRNAseq) allows unbiased and significantly more in-depth profiling of immune cell populations in animal models and humans in both healthy and diseased states. Because scRNAseq can define the transcriptomic heterogeneity of a complex community of cells and assign unbiased identity classifications to cell populations, it is optimally suited for the study of complex inflammatory states such as the one engendered by SARS-CoV-2 infection. ScRNAseq has recently identified initial cellular targets of SARS-CoV-2 infection in model organisms ^6-8^ and patients ^9-13^ and characterized peripheral and local immune responses in severe COVID-19 ^14^, with severe disease being associated with a cytokine storm and increased neutrophil accumulation. However, the human studies have mostly been performed in peripheral blood samples^14^, BAL^9^ and tissues^8,15^ from a limited number of moderate or severe COVID-19 patients within limited age ranges. To overcome the limitations associated with longitudinal early immune profiling in human subjects and to get more in-depth understanding of the early dynamics of transcriptional changes during COVID-19, we characterized the transcriptional signatures at the single cell level in the broncho-alveolar compartment of rhesus macaques at pre-infection collected 7 days before infection (−7dpi), at early stage of SARS-CoV-2 infection (3dpi) and at endpoint of the study (14-17dpi). Thus, the immune landscape in the broncho-alveolar compartment of SARS-CoV-2 infected adult rhesus macaques serves as a surrogate of early immune dynamics of protective immune responses in lungs after SARS-CoV-2 infection. We observed the appearance of distinct macrophage and T-lymphocyte populations exhibiting IFN-driven inflammatory gene signatures at 3dpi. The IFN responsive macrophage populations upregulated ACE2 expression and were infected by SARS-CoV-2. Further analysis of upregulated genes in the macrophages revealed IFN-driven innate antiviral defense and negative regulation of viral genome replication, suggesting a prominent role of macrophages driven innate immunity in resolution of SARS-CoV-2 infection.

## RESULTS

### Landscape of immune cells in the BAL of rhesus macaques during SARS-CoV-2 infection and recovery

To understand the early immune responses generated by SARS-CoV-2 in NHP model of COVID-19, we analyzed cryopreserved single cells isolated from BAL of young rhesus macaques infected by SARS-CoV-2 ^4^ at the following time points: 7 days prior to infection (−7dpi), three days post infection (3dpi), endpoint (14-17dpi) (Fig 1a). These time points were validated to represent distinct phases of viral dynamics *in vivo* covering baseline, the peak of acute viral infection (3dpi) and at the time that infection had resolved (endpoint) (Fig 1b) ^4^.

**Figure 1.**
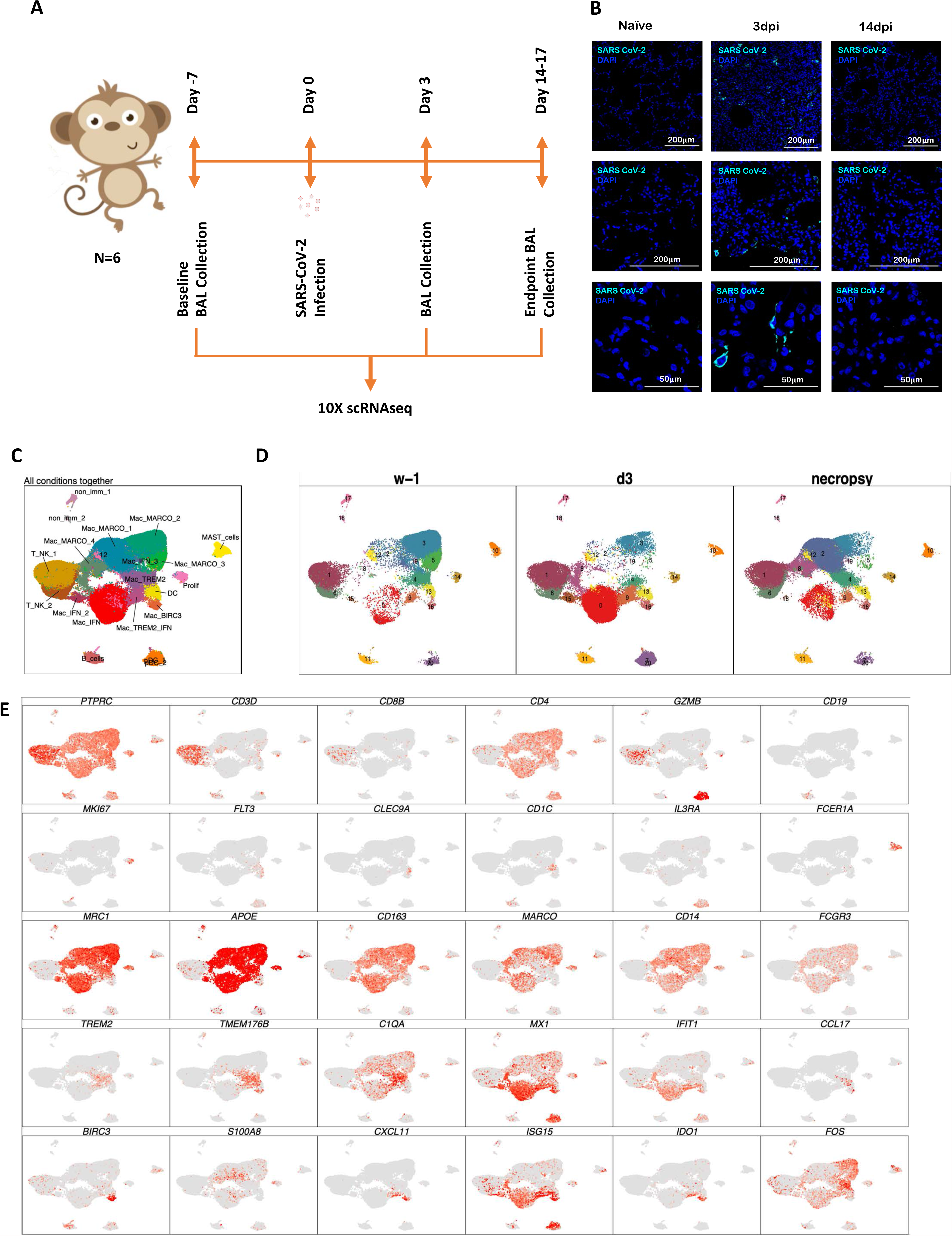
Immune landscape of BAL in SARS-CoV-2 infected macaques. Study outline of scRNAseq analysis of BAL cells from rhesus macaques infected with SARS-CoV-2. BAL single cell suspensions from 6 young rhesus macaques infected with SARS-CoV-2 from pre-infection (−7dpi), 3dpi and endpoint (14-17dpi) were subjected to scRNAseq (A). Immunofluorescence confocal images of the lungs stained with nucleocapsid (N)-specific antibodies (turquoise) and 4,6-diamidino-2-phenylindole (DAPI) (blue). Shown are the images at 10x, 20x and 63x magnification from Naïve lungs (uninfected) as well as lungs infected with SARS-CoV-2 at Day 3 and Day 14 post infection (B). UMAP plots of cells from all scRNAseq samples together, colored according to cluster classification (C) or respective timepoints (D). (E) UMAP plots with the expression of markers, characterizing main immune populations. n = 6.

We subjected single cells isolated from BAL of the SARS-CoV-2 infected rhesus macaques to 3’ 10x Genomics based scRNAseq processing and analysis pipeline with rigorous QC threshold (Fig S1) at -7dpi (n = 6), 3dpi (n = 6), and 14-17dpi (n = 6). Sequencing yielded a total of 170078 cells ranging from 1543-16608 cells per sample. The mean number of cells per sample in -7dpi was 8484, 3dpi was 9840 and 14-17dpi was 10021. Majority of the cells were immunocytes (Fig 1c) distributed across all time points (Fig 1d) and animals (Fig 1c, Fig S2). Consistent with the prior reports on cellular composition of BAL in rhesus macaques, myeloid were abundant comprising 77% of total cell than lymphoid compartment with 22% cells across all time points. The populations were homogenously distributed across all animals and across timepoints (Fig S2b). We identified 19 distinct cell clusters spanning varied cell types based on expression of canonical genes-T cells: Cluster of differentiation (CD) 3D^+^; Natural Killer (NK) cells: Killer Cell Lectin Like Receptor C3(KLRC3)^+^, Granzyme B (GZMB)^+^; B cells: CD19^+^, CD20/ Membrane Spanning 4-Domains A1 (MS4A1)^+^, CD79A^+^; macrophages: CD68^+^, CD163^+^) Dendritic Cells (DC): fms-like tyrosine kinase 3 (FLT3)^+^; conventional DCs (cDC): CD1c^+^; plasmacytoid DCs (pDC): (CD123/Interleukin (IL) 3 Receptor Subunit Alpha (IL3RA)^+^, CD303/ C-Type Lectin Domain Family 4 Member C (CLEC4C)^+^, Leukocyte Immunoglobulin Like Receptor A4 (LILRA4)^+^ and mast cells: (CD117/ KIT Proto-Oncogene (KIT)^+^, Fc Fragment Of IgE Receptor 1a (FCER1A)^+^, Carboxypeptidase A3 (CPA3)^+^ (Fig 1c,e, Fig 2a). Day 3 was distinctively marked by appearance of cell population expressing distinct IFN responsive gene signature comprising of MX Dynamin Like GTPase (MX) 1, MX2, Interferon Induced Protein With Tetratricopeptide Repeats (IFIT) 1, IFIT2, IFIT3, IFIT5, IFN-Inducible Protein ^16^ 6, IFI16, IFI44, Interferon-stimulated gene (ISG) 15, HECT And RLD Domain Containing E3 Ubiquitin Protein Ligase (HERC) 5, Sialic Acid Binding Ig Like Lectin (SIGLEC) 1, 2’-5’-Oligoadenylate Synthetase (OAS) 1, OAS2, OAS3 (Figure 2b). Although IFN-α and ACE2 transcripts were not abundantly present in the scRNAseq dataset, confocal analysis validated strong upregulation of IFN-α and ACE-2 in lungs of macaques on 3dpi compared to healthy or 14-17dpi (Fig 2c,d). Confocal analyses also validated higher expression IFN responsive transcripts like MX1 (Fig2e), MX2 (Fig2f) and ISG15 (Fig2g) in lung tissues isolated from rhesus macaques at 3dpi (Fig 2c) when compared to 14-17dpi and healthy macaques.

**Figure 2.**
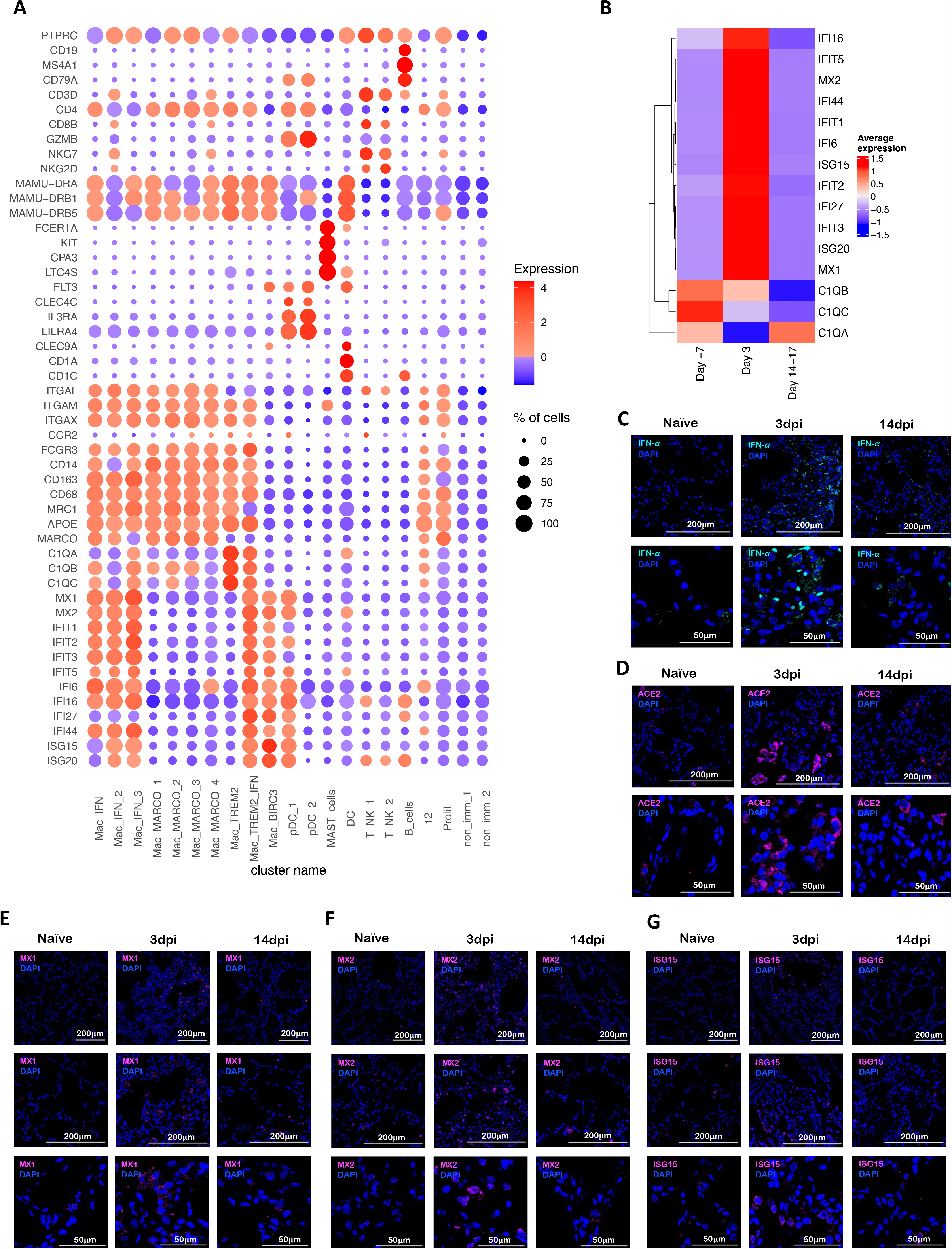
SARS-CoV-2 infection induces IFN responsive gene signature in rhesus macaques. (A) Bubble plot showing the fold change of genes in identified cell clusters and the fraction of cells expressing the gene of interest. (B) Heatmap of key interferon responsive genes at different timepoints. Confocal images validating *in vivo* expression of IFN-α (turquoise) (C), ACE2 (magenta) (D), MX1 (magenta) (E), MX2 (magenta) (F) and ISG15 (magenta) (G) with DAPI (blue) in the lung sections of Naïve rhesus macaques and SARS-CoV-2 infected lungs at Day 3 and Day 14 post infection.

As described previously, in BAL, SARS-CoV-2 vRNA levels were detected in of 5/6 macaques at 3dpi by RT-qPCR. Virtually no BAL vRNA was detected at the endpoint suggesting that the rhesus macaques cleared the virus, from the BAL compartment ^4^. vRNA in Nasal Swabs (NS) could be detected in only 4 animals at Day 3, while all animals recorded vRNA at Day 9, and only 3 at the endpoints. vRNA was detected in the lungs of 3 macaques at necropsy (14-17dpi) while no SARS CoV-2 subgenomic RNA (correlate for infectious/replicating virus) was detected in any rhesus macaque lungs at Endpoints ^4^. No vRNA was detected in any plasma samples or in randomly selected urine samples. Based on vRNA persistence in the lungs of immunocompetent young macaques and the absence of replicative virus, it was concluded that that macaques efficiently control SARS-CoV-2 infection in a duration of two weeks ^4^. In lieu of the absence of replicating virus in lungs at endpoint and peak viremia in 3dpi BAL, in vivo pathology was also found to peak around 3dpi and subsided thereafter as shown by chest x-ray scores. Furthermore, tissue pathologies were observed in lungs at endpoints suggesting viral antigen triggered immune response by persisting antigen. Along with the absence of replicating virus in lungs at endpoint pathological observations were found as shown by histological analyses endpoint ^4^. Our previous immunological analyses of BAL cells by flow cytometry had revealed a massive infiltration of immunocytes mainly comprising of T cells, interstitial macrophages, neutrophils and plasmacytoid Dendritic Cells ^4^. Appearance of these populations also correlated strongly with the viral loads ^4^. When combined with these previously reported findings, our new results clearly establish influx of myeloid cells and induction of a fully functional IFN driven innate immune response in macrophages against SARS-CoV-2 in the lungs of rhesus macaques. Our scRNA-seq based deep cellular phenotyping analysis clearly establishes the induction of a robust and targeted innate immune response mainly driven by macrophages as opposed to dysregulated immune responses against SARS-CoV-2 in the early phase of infection.

### Myeloid bronchoalveolar landscape

A total of 129280 myeloid cells were analyzed across all timepoints which distributed into 17 distinct clusters across the 3 timepoints studied (Fig 3A-B, Fig S3a). The populations were homogenously distributed across all animals at all timepoints (Fig S3b). We noted distinctive cluster alignment of all myeloid populations based on key myeloid phenotype markers (Fig 3c-d) that differed between different conditions (Figure 3d).

**Figure 3.**
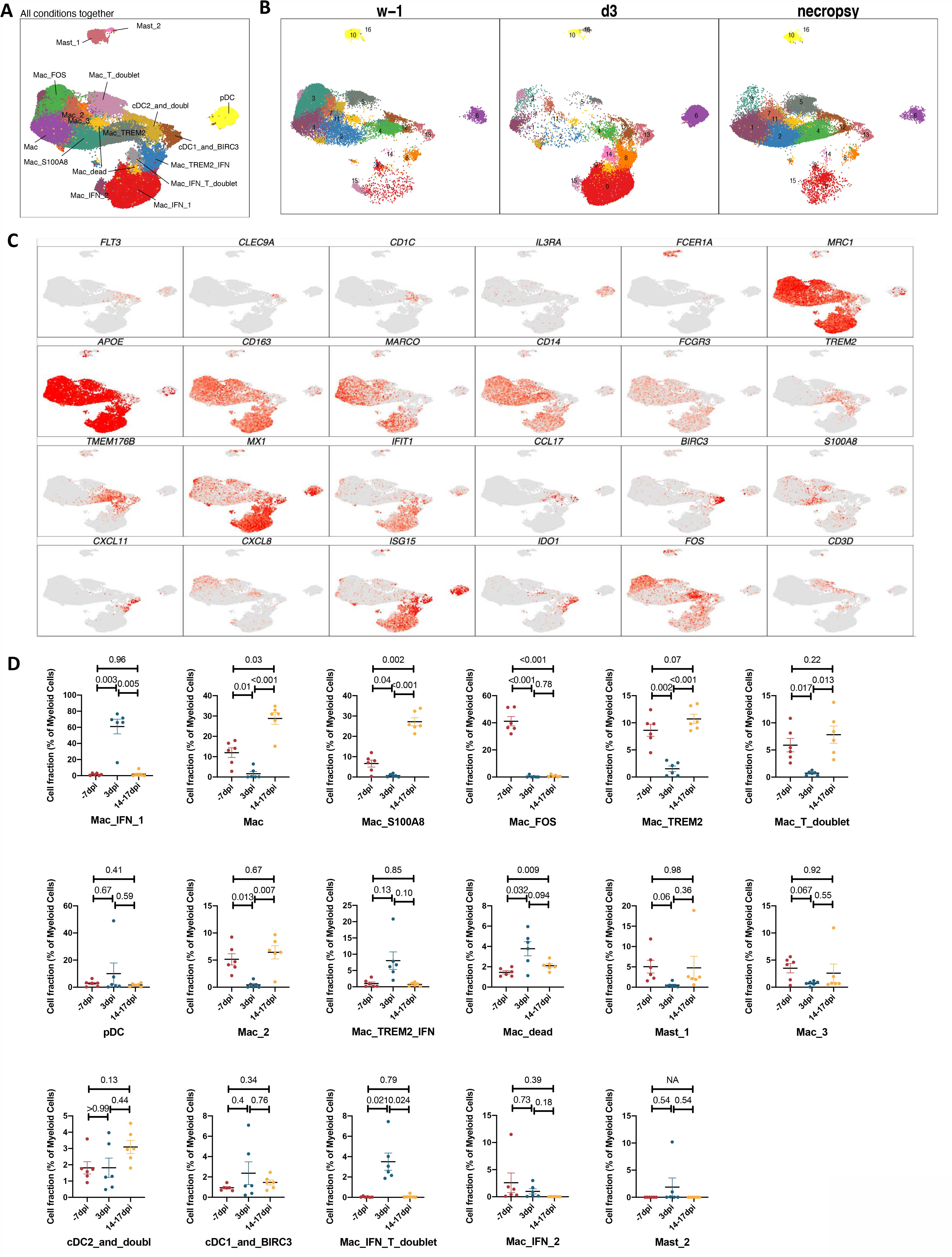
Myeloid single cell landscape in SARS-CoV-2 infected macaques. BAL myeloid cell dynamics in macaques infected with SARS-CoV-2 by scRNA-seq demonstrates the presence of IFN response in pDCs and IFN-responsive macrophage compartments. (A) UMAP plot of myeloid cells from all scRNA-seq samples together, colored according to (A) cluster classification or respective (B) timepoints. (C) UMAP plots with the expression of markers, characterizing main myeloid populations in macaques. (D) Cell proportion of each cluster per condition. n = 6.

As expected, due to limitation of the 10x Genomics scRNAseq pipeline to detect neutrophils^17^, this population was not detected in our analysis. As reported earlier in various NHP studies ^4,18,19^, BAL landscape mostly comprised of macrophages, which are distributed into alveolar (CD206^+^) or interstitial (CD206^-^) phenotypes ^4,19^. Our prior scRNAseq analysis using the 10x platform in single cells isolated from lungs of rhesus macaques with tuberculosis had identified novel macrophage phenotypes exhibiting distinct TREM2 and IFN-responsive gene signatures^17^. Here, we found 3 distinct IFN-responsive macrophages populations which were predominantly present on 3dpi (Fig 4a,b), one of which also expressed high levels of Triggering Receptor Expressed On Myeloid Cells (TREM) 2 gene expression (Fig 4a). For reference, we annotated the most abundant IFN-responsive macrophage population as Mac_IFN_1, second IFN responsive macrophage population was annotated Mac_IFN_2 and the third IFN responsive macrophages with a TREM2 expression module were annotated Mac_TREM2_IFN (Fig 4a). pDCs are considered the chief drivers and source of Type I IFN signature. Our prior data suggested a significant influx of pDCs in BAL compartment and lungs of SARS-CoV-2 infected macaques^4^. The pDC cluster in our current scRNAseq data was identified by expression of classic pDC markers like IL3RA/CD123, CLEC4C and Transcription Factor (TCF) 4. This cluster was only modestly increased at 3dpi. However, the pDC cluster contained genes associated with innate response to viral pathogens like Toll-like receptors (TLR)7, TLR9 etc. along with induction of Type I IFN response like Interferon Regulatory Factor (IRF) 1, IRF3, IRF7, IRF8, IRF9, Derlin (DERL) 3, Solute Carrier Family 15 Member 4 (SLC15A4), with significant induction in expression levels. In addition, expression of an IFN responsive transcriptional signature (MX1, MX2, ISG15, ISG20, IFI6, IFI16, IFI27) was also significantly elevated in this pDC cluster (Fig 4c, d). Multilabel confocal microscopy analysis validated the higher expression of IFN-α by pDCs in lung tissues isolated from infected macaques at 3dpi (Fig 4e) when compared to 14-17dpi and healthy macaques (Fig4e, S4).

**Figure 4.**
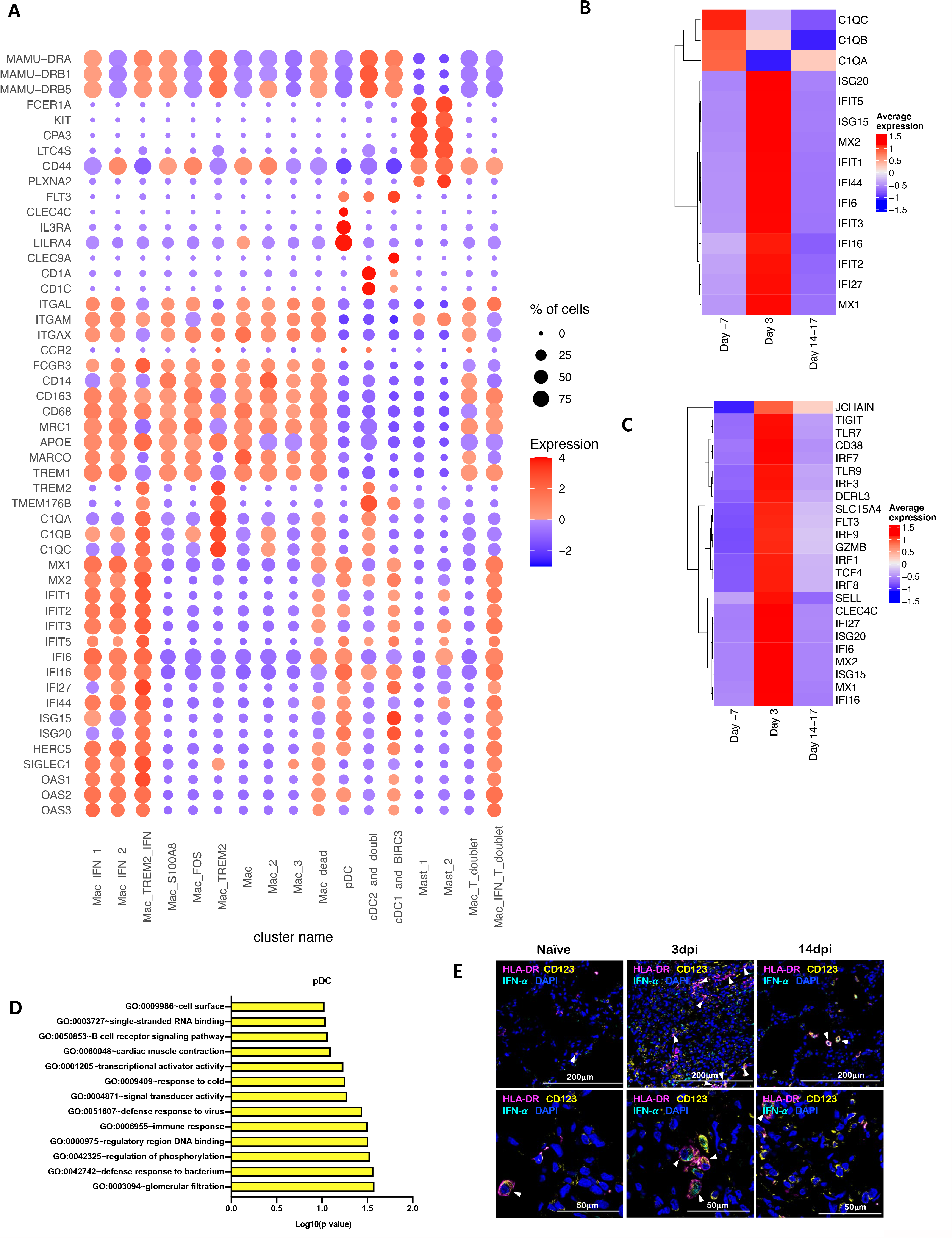
Macrophages and pDCs are the dominant cells driving Type I IFN response in lungs of SARS-CoV-2 infected macaques. (A) Bubble plot showing the fold change of genes in identified myeloid cell clusters and the fraction of cells expressing the gene of interest. (B) Heatmap of key interferon responsive genes at different timepoints in macrophage clusters. (C) Heatmap of key interferon responsive genes at different timepoints in plasmacytoid dendritic cells sub-clusters. (D) GO pathways enriched in upregulated genes in pDCs. (E) Multilabel immunofluorescence confocal images validating *in vivo* expression of IFN-(turquoise) in pDCs marked by HLA-DR (magenta) and CD123 (yellow) in Naïve Rhesus macaque lungs as well as at Day 3 and Day 14 post infection with SARS CoV-2. Shown are the images at 20x and 63x magnification.

Among macrophage subclusters, mac_IFN_1 was the most abundant population found in 3dpi BAL samples, comprising 70 percent of myeloid cells; this population was completely absent in pre-infection and during resolution at endpoint. IFN-responsive gene signature was strongly upregulated in this population and the key genes which were most differentially upregulated in this population were MX1, MX2, IFIT1, IFIT2, IFIT3, IFIT5, IFI6, IFI16, IFI44, ISG15, HERC5, SIGLEC1, OAS1, OAS2, OAS3 etc. (Fig 4a). MX1 encodes a guanosine triphosphate (GTP)-metabolizing protein called IFN-induced GTP-binding protein Mx1 which is induced by Type I and Type II IFNs, antagonizes the replication process of several RNA and DNA viruses and participates in the cellular antiviral responses ^20-23^. MX2, a paralog of MX1, is another IFN-induced GTP-binding protein that induces innate antiviral immune responses. IFIT genes encode IFN-induced antiviral proteins which acts as an inhibitor of cellular as well as viral processes, cell migration, proliferation, signaling, and viral replication^24,25^. IFI6 is one of the earliest identified IFN induced gene encoding the IFN-α -inducible protein 6 which has been shown to exert antiviral activity towards viruses by inhibiting the EGFR signaling pathway^26-29^. IFI16 gene encodes the Interferon Gamma Inducible Protein 16 which has been shown to be involved in the sensing of intracellular DNA and inducing death of virus infected cells^30-33^. IFI44 encodes the Interferon-Induced Protein 44 which is induced by Type 1 but not Type II IFNs and has been reported to suppress viral transcription ^34,35^. IFIT genes encodes for the IFIT proteins (Interferon Induced proteins with Tetratricopeptide repeats) which confer antiviral state in a cell by either directly binding to the viral RNA or by binding to eukaryotic initiation factor 3 (eIF3) and preventing eIF3 from initiating viral translational processes ^36-38^. All four classes of IFIT i.e. IFIT1, IFIT2, IFIT3 and IFIT5 were upregulated in the mac_IFN_1 population (Fig 4a). ISG15 also called Ubiquitin-like protein ISG15 is an early mediator of signaling induced by Type I IFNs and elicits innate immune response to viral infections by conjugation/ISGylation of its targets like MX and IFIT ^39-42^. OAS encodes IFN-induced, dsRNA-activated antiviral enzyme which plays a critical role in cellular innate antiviral responses ^43,44^. HERC5 is an E3 ligase for ISG15 conjugation which acts as a positive regulator of innate antiviral response in cells induced by IFNs and functions as part of the ISGylation machinery ^45-47^. SIGLEC1 (CD169) is an IFN-inducible gene that acts as an endocytic receptor mediating clathrin dependent endocytosis and has been reported to be upregulated in circulating monocytes in COVID-19 patients ^48-51^. Gene set enrichment analysis (GSEA) analysis revealed defense response to virus, negative regulation of viral genome replication, response to IFN-α and innate immune response as enriched gene ontology (GO) terms in this population (Fig 5a).

**Figure 5.**
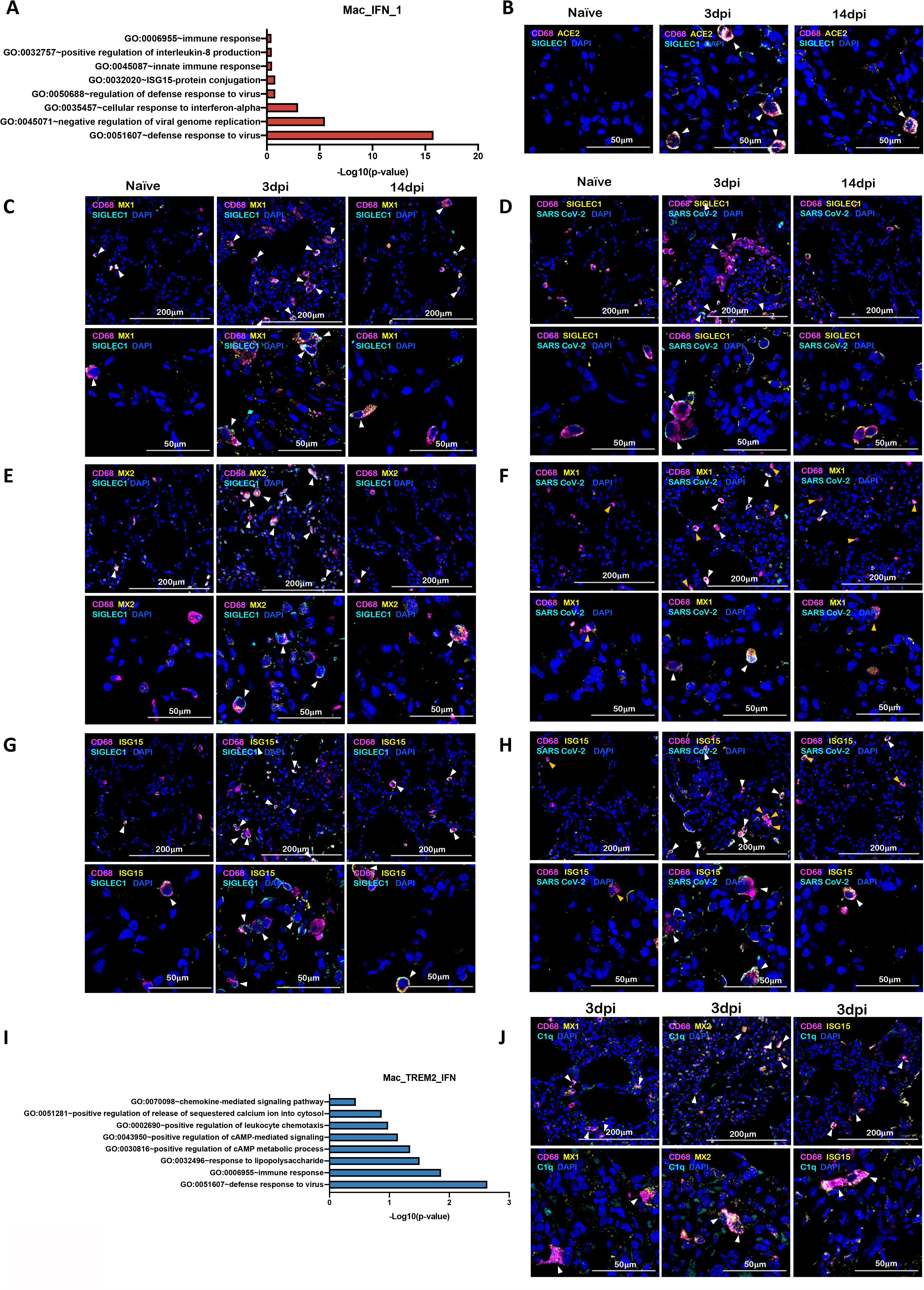
IFN induced viral defense response in lung macrophages of SARS-CoV-2 infected macaques. (A) GO pathways enriched in upregulated genes in Mac_IFN_1 subcluster. Multilabel immunofluorescence confocal images validating *in vivo* expression of (B) ACE2 (yellow) and SIGLEC1 (turquoise), and (C) MX1 (yellow) and SIGLEC1 (turquoise) in macrophages (magenta), (D) CD68 (magenta) and SIGLEC1 (yellow) positive macrophages harboring SARS CoV-2 (turquoise), (E) Macrophages (magenta) expressing MX2 (yellow) and SIGLEC1 (turquoise), (F) MX1 (yellow) positive macrophages (magenta) with SARS CoV-2 (turquoise), (G) Macrophages (magenta) expressing ISG15 (yellow) and SIGLEC1 (turquoise), (H) SARS CoV-2 (turquoise) harbored in ISG15 (yellow) expressing macrophages (magenta) in lungs of Naïve and 3 and 14 days post infection of SARS CoV-2 infected macaques. Nuclei stained with DAPI are shown in blue. White arrows represent macrophages expressing ACE2 and SIGLEC1 in (B); MX1 and SIGLEC1 in (C); MX2 and SIGLEC1 in (E); ISG15 and SIGLEC1 in (G). In (D), (F) and (H), white arrows mark the presence of SARS CoV-2 in macrophages expressing SIGLEC1, MX1 and ISG15 respectively; whereas, orange arrows are used to mark SIGLEC1, MX1 and ISG15 expressing macrophages with no SARS CoV-2. (I) GO pathways enriched in upregulated genes in Mac_TREM2_IFN subcluster. (J) Multilabel confocal immunofluorescence images validating *in vivo* expression of MX1, MX2 and ISG15 (shown in yellow) in TREM2 macrophages in lungs of SARS-CoV-2 infected macaques at 3dpi.

SARS-CoV-2 infection in CD169^+^ macrophages has been reported in COVID-19 ^15,52^. To validate our findings of SARS-CoV-2 infection in CD169^+^ macrophages, lung tissues from rhesus macaques following SARS CoV-2 infection and healthy controls were stained for CD68, ACE2, SIGLEC1, MX1, MX2, ISG15, Complement component 1q (C1q) and SARS CoV-2 nucleocapsid antibody (Fig5 b-h, j, S5-12). Lung macrophages expressing SIGLEC1/CD169 were enriched and expressed high levels of ACE2 at 3dpi (Fig 5b, S5). Another independent study has established that human macrophages and monocytes can be infected by SARS-CoV-2 but that the infection is abortive^53^. The macrophage population was further studied for detecting IFN responsive elements: MX1 (Fig 5c,S6), MX2 (Fig 5e, S8), ISG15 (Fig 5g, S10) and viral antigens (Fig 5d, S7, 5f, S9, 5h, S11) and were found to be abundantly harboring SARS-CoV-2 *in vivo* as shown by multicolor confocal staining for SARS-CoV-2 Nucleocapsid/Spike protein in lung sections. MX1/MX2/ISG15 staining with viral antigen clarified that the IFN-responsive signature was mostly restricted to macrophages harboring SARS-CoV-2, confirming an early IFN-driven innate immune response in lung macrophages.

Mac_IFN_2 cluster was mostly identical to Mac_IFN_1 but expressed a comparatively higher expression of C-X-C Motif Chemokine Ligand. (CXCL) 8, IL1B and tumour necrosis factor (TNF) associated with nuclear factor-kappa B (NFKB) Inhibitor Zeta (NFKBIZ), NFKBIA, TNF-α-induced protein 3 (TNFAIP3) and Activator protein (AP)1 signaling: Fos Proto-Oncogene ^54^, Jun Proto-Oncogene (JUN), JunB Proto-Oncogene (JUNB).

We previously reported a novel cell cluster of alveolar macrophages abundantly found in lungs of rhesus macaques showing an enriched expression of TREM2, transmembrane protein (TMEM) 176A/B and C1Q genes ^17^. Our current data validated the presence of two clusters of macrophages showing TREM2 gene signatures, one of which expressed a strong IFN-β responsive gene signature. Therefore, we annotated them as Mac_TREM2 and Mac_TREM2_IFN respectively. Mac_TREM2 cluster was abundantly present in the BAL from healthy macaques and switched to an IFN-responsive phenotype on 3dpi which was then restored at the endpoint (Fig 3d). Mac_TREM2 cluster observed at pre-infection baseline and then replenished to normal levels at endpoint (Fig 3d) demonstrated enriched expression for FOS, FosB Proto-Oncogene (FOSB), Activating transcription factor (ATF) 3, Regulator Of G Protein Signaling ^55^ 1, Aryl hydrocarbon receptor (AHR), NFKBIZ, BTG Anti-Proliferation Factor (BTG) 2, Early Growth Response (EGR) 1, Lamin A/C (LMNA), RasGEF Domain Family Member 1B (RASGEF) 1B and CD69 (Fig 4a). While the Mac_IFN_TREM2 cluster showed an upregulation of IFN-responsive gene signature comprising of SIGLEC1, MX, IFIT, IFI, ISG and OAS genes (Fig 4a), suggesting defense response to virus as significantly enriched geneset (Fig5i). *In vivo* validation in lungs of macaques confirmed abundance of TREM2 macrophages with IFN-responsive phenotype on 3dpi (Fig 5j, S12).\ Mac_FOS cluster was abundant in BAL at baseline and constituted ∼40-50% of myeloid cells. However, this cluster was depleted at 3dpi and not restored at the endpoint tested. Mac_FOS expressed high CXCL8 expression along with FOS, FOSB, NFKBIA, NFKBIZ AHR, lysozyme (LYZ) and CD69. GSEA revealed that this subset expressed an inflammatory and neutrophil chemotactic nature (Fig S13a), even in absence of infection in healthy macaques.

Mac_S100A8 cluster constituted 10% of the myeloid cells at pre-infection baseline, but was absent on 3dpi and abundantly represented at the endpoint suggesting a potential role for these cells in SARS-CoV-2 mediated pathology (Fig 3d). The key genes upregulated in this cluster were S100 Calcium Binding Protein (S100) A4, S100A6, S100A8, S100A9, Cathelicidin Antimicrobial Peptide (CAMP), Carboxypeptidase Vitellogenic Like (CPVL) (Fig4a) which represent innate inflammatory immune responses, neutrophil aggregation and chemotaxis pathways (Fig S13b). Mac_S100A8 cluster has four genes significantly upregulated from the S100 family of genes that involves low molecular-weight proteins considered as potent damage-associated molecular pattern molecules (DAMPs). DAMPs are also called danger signals or alarmins as they serve as a warning sign for the innate immune system to alert ambient damage or infection. S100A8 protein also called calgranulin A forms a heterodimer with S100A9 protein called calgranulin B, to form a heterodimer called Calprotectin which stimulates T-lymphocyte chemotaxis by acting as a chemoattractant complex with Peptidoglycan Recognition Protein 1 (PGLYRP1) that promotes lymphocyte migration via C-C chemokine receptor (CCR) 5/ C-X-C motif chemokine receptor (CXCR) 3 receptors^56,57^; neutrophil recruitment along with TLR4 and/or receptor for advanced glycation end products ^58^ -mediated multiple inflammatory pathways ^59,60^. Intracellular functions of S100A6 includes regulation of several cellular functions, such as proliferation, apoptosis, cytoskeleton dynamics, response to different stress factors etc. but when secreted into extracellular milieu it also RAGE (receptor for advanced glycation end products) and integrin β1 mediated inflammatory responses^61^. S100A4 has been reported to synergize with vascular endothelial growth factor (VEGF) in a RAGE dependent manner to promoting endothelial cell migration by increasing KDR (kinase insert domain receptor)/Vascular Endothelial Growth Factor Receptor 2 (VEGFR2) expression and MMP-9 activity^62^. S100A4 also plays a major role in high-density collagen deposition ^63^.

Three other populations of macrophages lacking IFN signature (annotated Mac, Mac_2 & Mac_3) are present in healthy macaques at pre-infection timepoint. These populations were non-existent at 3dpi but are replenished at the endpoints (Fig 3d, 4a).

To further understand the influence of macrophages on ambient immunocytes, we analyzed the ligand-receptor interactions between the most abundant macrophage population and other immunocytes based on cell specific transcripts read at respective timepoint. The ligand-receptor interactions between Mac_IFN_1 and other immunocytes present at 3dpi depicted as Circos plot (Fig 6a) shows a prominent co-stimulatory potential of Mac_IFN_1 population on Mast cells via Adrenoceptor Beta (ADRB) 2, SIGLEC10, Cholinergic Receptor Muscarinic (CHRM) 3; cDCs via LDL Receptor Related Protein (LRP) 2, TNF Receptor Superfamily Member (TNFRSF) 11B, and pDCs via Nerve Growth Factor Receptor (NGFR), CD28. Mac_FOS showed prominent interaction with cDCs (Fig 6b) and the Mac_S100A8 cluster with Mast cells, cDCs, pDCs in addition to Mac_TREM2_IFN (Fig 6c).

**Figure 6.**
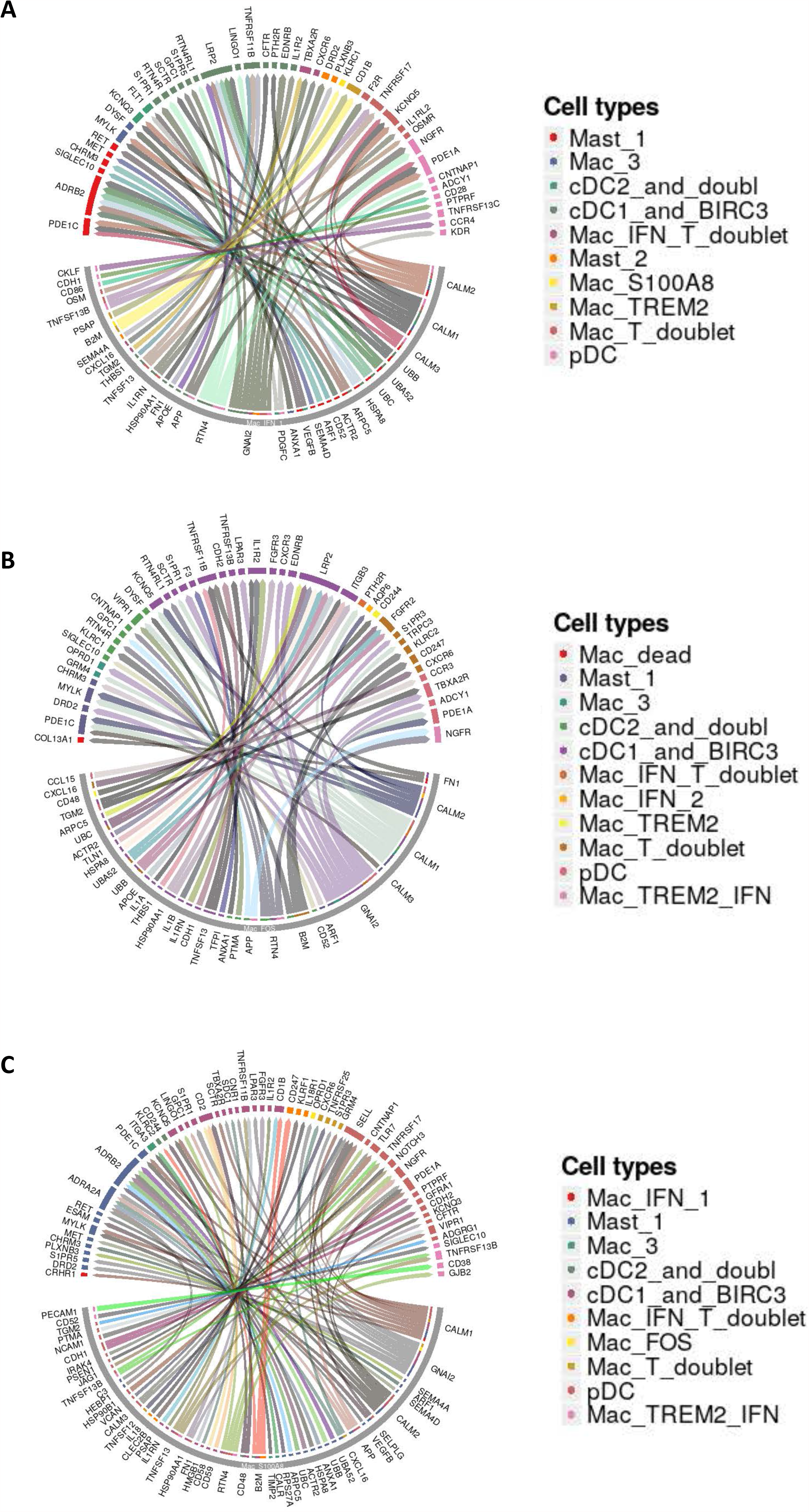
Macrophage interactome model in SARS-CoV-2 infected lungs. Macrophage-Immunocytes interactome. Circos plots showing the ligand-receptor interactions between most abundant macrophage populations and different immune cell in the three conditions studies. (A) Circos plot depicting interaction of Mac_IFN_1 with ambient immunocytes based on ligand-receptor transcript reads at 3dpi. (B) Circos plot depicting interaction of Mac_S100A8 with ambient immunocytes based on ligand-receptor transcript reads at 14-17dpi. (C) Circos plot depicting interaction of Mac_FOS with ambient immunocytes based on ligand-receptor transcript reads at -7dpi.

### Lymphoid bronchoalveolar landscape

A total of 38160 lymphoid cells were analyzed across all timepoints which distributed into 13 distinct clusters across the 3 timepoints studied (Fig 7a, Fig S14a,b). The populations were homogenously distributed across all animals at each timepoint (Fig S14c). We noted distinctive cluster alignment of all lymphoid populations based on key lymphocyte phenotype markers (Fig 7a,b) that differed between different conditions (Figure 7c,d). The only distinct lymphocyte cluster found to be upregulated at Day 3 post infection was a T cell cluster with IFN responsive gene signature spanning MX1, MX2, ISG15, ISG20, IFI27, IFI44, IFIT1, IFIT2, IFIT3, IFIT5, OAS1, OAS2, HERC5 HERC6 genes and was annotated T_IFN (Fig 7d,e). Confocal analysis in lung sections validated the abundance of IFN responsive T cells at 3dpi in macaque lungs (Fig7 f, S15).

**Figure 7.**
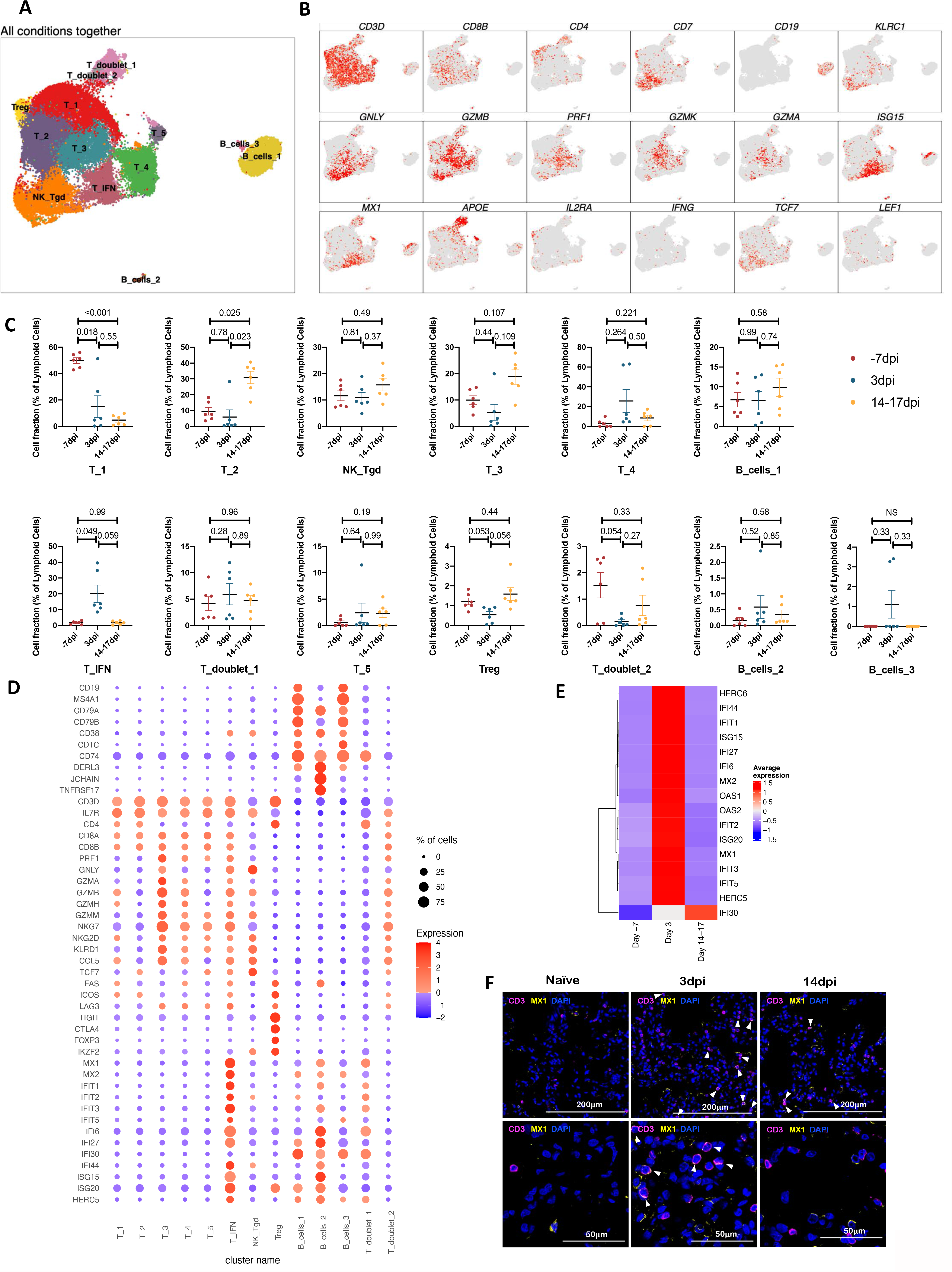
Lymphoid single cell landscape in SARS-CoV 2 infected macaques. BAL lymphoid cell dynamics in macaques infected with SARS-CoV-2 by scRNA-seq demonstrates the presence of IFN responsive T cells. (A) UMAP plot of lymphoid cells from all scRNA-seq samples together, colored according to cluster classification. (B) UMAP plots with the expression of markers, characterizing main lymphoid populations in macaques. (C) Cell proportion of each cluster per condition. n = 6. (D) Bubble plot showing the fold change of genes in identified lymphoid cell clusters and the fraction of cells expressing the gene of interest. (E) Heatmap of key interferon responsive genes at different timepoints in lymphoid sub clusters. (F) Multilabel immunofluorescence confocal images validating *in vivo* expression of MX1 (yellow) in T-cells marked by CD3 (magenta) and nuclei (blue) in Naïve as well as SARS CoV-2 infected lungs at 3dpi and 14dpi. White arrows represent T cells expressing MX1.

Regulatory T cells (T_Regs_) constituted less than 3% of the lymphoid population and expressing negative checkpoint regulators (NCR) like inducible costimulator (ICOS), Lymphocyte Activating (LAG) 3, T Cell Immunoreceptor With Ig And ITIM Domains (TIGIT) etc. along with forkhead box P3 (FOXP3). When compared to pre-infection baseline T_Regs_ showed a slight decline at 3dpi but were replenished at the endpoints. IFN-α can drive contraction of T_Regs_ while ISG15 can rescue T_Regs_ from Type I IFN induced contraction ^64^. We observed high IFN-α concentrations at day 3, and relatively lower expression of ISG15. These results could explain the contraction of T_Regs_ ar 3 dpi.

## DISCUSSION

A thorough understanding of the host inflammatory responses during SARS-CoV-2 infection is needed both the identification of correlates of protection versus pathology. Such information is also critical in order to identify pathways which can be precisely modulated to limit inflammation without affecting protective mechanisms. Recognition of viral infections by innate immune sensors activates both the Type I and Type III IFN responses. Elevated levels of IFNs have been reported to correlate with and contribute to severe COVID-19^5,65-73^. It is possible that severe infection drives an uncontrolled expression of the Type I IFN response leading to pathology instead of viral containment. However, inborn errors of Type I IFN in COVID-19 patients have been associated with life threatening conditions^67,68^. Similar life threatening manifestation of COVID-19 has also been attributed to patients having autoantibodies against type I IFNs^73^. It is possible that delayed or inadequate IFN responses lead to inflammation mediated damage during later phases of disease. Increased induction of early Type I IFN signaling pathways in SARS-CoV-2 infected macaques suggest a role for IFN signaling in protection rather than disease progression^4^. This is further supported by the increased induction of Type I IFN signaling in the cohort of young relative to geriatric macaques^5^, suggesting that IFN induction may be compromised in older or immunocompromised hosts^5,74^. Thus, it is not fully clear if Type I IFNs are protective or pathological in COVID-19 ^74^. This is an important paradox to resolve, as it could lead to better therapeutic approaches for COVID-19 as well as for long-term persistent COVID-19 sequelae. Elegant approaches are available to modulate the signaling of this pathway in macaques^75^. Our studies lay the foundation of Type I IFN depletion studies in this model to better understand the role of this pathway in the early control of SARS-CoV-2 infection and in limiting inflammation. Further testing the protective versus pathological roles of IFNs in different phases of COVID-19 in the macaque model with the availability of IFNAR blocking reagents should further clarify the specific role of IFN pathways in COVID-19.

Our results unequivocally show that in protected, immunocompetent hosts, SARS-CoV-2 infection is characterized by an acute inflammatory response leading to a myeloid cell influx into the lung compartment^4^ and a strong Type I IFN response^5^. Using state-of-the-art scRNAseq approach in longitudinal BAL samples, we now demonstrate that the robust Type I IFN and related cytokine response observed in the lungs of infected macaques is mediated by myeloid cell subpopulations that are alveolar rather than tissue-resident in nature. In particular, macrophage subpopulations Mac_IFN_1, Mac_IFN_2 and Mac_TREM2_IFN subpopulations expressed high levels of IFN downstream genes both in magnitude and frequency (Fig 2). Our results clearly show that induction of a robust IFN response in macrophages strongly correlates with viremia (Fig S16a,b) and subsequent clearance of SARS-CoV-2 from the airways of macaques (Fig S16c,d).

## Supplement

Legends to Supplements:

**Fig S1**. UMAP plots representing QC parameters for analysis. (A) nUMI, (B) nGenes and (C) fraction of mitochondrial genes.

**Fig S2**. (A) UMAP plots depicting all clusters (left), cluster annotations (middle), distribution as per condition (right) and (B) distribution per sample.

**Fig S3**. (A) UMAP plots depicting all myeloid clusters (left), cluster annotations (middle), distribution as per condition (right) and (B) distribution per sample.

**Figure S4**: Single channel images depicting pDCs marked with HLA-DR (magenta) and CD123 (yellow) showing expression of IFN-α (turquoise) in Naïve lungs as well as SARS CoV-2 infected lungs at Day 3 and Day 14 post infection at 63x magnification. Nuclei stained with DAPI are shown in blue.

**Figure S5**: Single channel images validating the expression of ACE2 receptor (yellow) on CD68 (magenta) and SIGLEC1 (turquoise) positive macrophages in Naïve and SARS CoV-2 infected lungs (3dpi and 14dpi).

**Figure S6**: Images showing separate channels for expression of Mac-IFN signature marker MX1 (yellow) by CD68 (magenta) and SIGLEC1 (turquoise) positive macrophages in Naïve as well as 3dpi and 14dpi SARS CoV-2 infected macaque lungs.

**Figure S7**: Single color images from Naïve and SARS CoV-2 infected lungs (3dpi and 14dpi) depicting SARS CoV-2 (turquoise) is harbored by CD68 (magenta) and SIGLEC1 (yellow) positive macrophages.

**Figure S8**: Shown are the images for single channels for CD68 (magenta) and SIGLEC1 (turquoise) positive macrophages expressing MX2 (yellow), one of the Mac-IFN signature marker. Images from Naïve as well as SARS CoV-2 infected lungs at Day 3 and Day 14 post infection were taken to see the differential expression.

**Figure S9**: Shown are the single channel images from Naïve and SARS CoV-2 infected lung sections (3dpi and 14dpi) depicting CD68 (magenta) and MX1 (yellow) positive macrophages harbors SARS CoV-2 (turquoise).

**Figure S10**: Single color images from the Naïve and SARS CoV-2 infected lungs sections (at 3dpi and 14dpi) showing CD68 (magenta) and SIGLEC1 (turquoise) positive macrophages expressing ISG15 (yellow), another Mac-IFN signature marker.

**Figure S11**: Single color images depicting ISG15 (yellow) expressing macrophages (magenta) harbor SARS CoV-2 (turquoise). Images shown are from the Naïve and SARS CoV-2 infected lung sections both at 3dpi and 14dpi.

**Figure S12**: Single channel images from lungs of SARS CoV-2 infected macaques at 3dpi showing the expression of Mac-IFN signature markers: MX1, MX2 and ISG15 (yellow) and TREM marker: C1q (turquoise) in CD68 positive macrophages (magenta).

**Fig S13**. GSEA enrichment set in (A) Mac_FOS and (B) Mac_S100A8 cluster.

**Fig S14**. (A) UMAP plots depicting all lymphoid clusters (left), cluster annotations (middle), distribution as per condition (right), (B) respective timepoints. and (C) distribution per sample.

**Figure S15**: Images showing single channels from CD3 (magenta) and MX1 (yellow) stained lungs sections from Naïve and SARS CoV-2 infected macaques at Day 3 and Day 14 post infection.

**Figure S16**: (A) Correlation curves depicting trends between fraction of cellular subsets and Log_10_ viral titers in BAL at the same time point. (B) Plot depicting Spearman correlation coefficient and associated p values. (C) Correlation curves depicting trends between fraction of cellular subsets and Log_10_ viral titers in BAL at the subsequent time point. (D) Plot depicting Spearman correlation coefficient and associated p values.

**Table S1**: List of animals and demographics.

**Table S2**: Details of antibodies.

## MATERIALS AND METHODS

### Macaques

No live Indian origin rhesus macaques were used in this study. Samples obtained from young Rhesus macaques *(Macaca mulatta)* infected with 1.05×10^6^ pfu SARS-CoV-2 isolate USA-WA1/2020 (BEI Resources, NR-52281, Manassas, VA) enrolled in a previously described (approved by the Animal Care and Use Committee of the Texas Biomedical Research Institute) study ^4^ were used for further analysis (Table S1).

### Isolation of BAL single cells from macaques

Single cell suspensions from BAL obtained at different time points were collected as described earlier ^4,76^ and cryopreserved in Cryostor-CS10 (Biolife Solutions, USA) at -70°C and then used for downstream processing of scRNAseq.

### Single cell RNA: Library generation and sequencing

scRNAseq was done according to the manufacturer instructions (10x genomics) and as previously described ^77^. Briefly, after quickly thawing the frozen BAL single cell suspension in water bath, 2×10^6^ cells were taken for downstream processing. BAL single cell suspensions were subjected to droplet-based massively parallel single-cell RNA sequencing using Chromium Single Cell 3’ (v3.1) Reagent Kit in the BSL-3 laboratory as per manufacturer’s instructions (10x Genomics). Briefly, cell suspensions were loaded at 1,000 cells/μL with the aim to capture 10,000 cells/lane. The 10x Chromium Controller generated GEM droplets, where each cell was labeled with a specific barcode, and each transcript labeled with a unique molecular identifier ^78^ during reverse transcription. The barcoded cDNA was isolated and removed from the BSL-3 space for library generation. The cDNA underwent 11 cycles of amplification, followed by fragmentation, end repair, A-tailing, adapter ligation, and sample index PCR as per the manufacturer’s instructions. Libraries were sequenced on a NovaSeq S4 (200 cycle) flow cell, targeting 30,000 read pairs/cell.

### Single cell RNAseq: data processing

The Cell Ranger Single-Cell Software 3.0 available at 10x website was used to perform sample demultiplexing. We aligned resulting fastq files on mmul10 genome (Genebank, https://www.ncbi.nlm.nih.gov/assembly/GCF_003339765.1/), with addition of Ensembl mmul8 mitochondrial genes for GTF file with cellranger count. For each sample the recovered-cells parameter was set to 10,000 cells that we expected to recover for each individual library.

We used R package Seurat 3 ^79^ for downstream analysis of count matrixes that we got as output from cellranger count^77^. We filtered cells that (1) had more than 10% of mitochondrial gene content and ^80^ had less than 363 detected genes. Data was log-normalized with a scale factor of 10^4^. The most variable genes were detected by the FindVariableFeatures function and used for subsequent analysis. Latent variables (number of UMI’s and mitochondrial content) were regressed out using a negative binomial model (function ScaleData). Principle component analysis ^55^ was performed with RunPCA function. A UMAP dimensionality reduction was performed on the scaled matrix (with most variable genes only) using the first 20 PCA components to obtain a two-dimensional representation of the cell states. For clustering, we used the functions FindNeighbors (20 PCA) and FindClusters (resolution 0.5). Identified clusters were split into 2 cell groups: myeloid and lymphoid - and ran through reclustering pipeline individually. For both cell subsets reclustering we performed clustering on first 20 PCA component. We used clustering resolution 0.35 for myeloid and lymphoid cells reclustering. To identify marker genes, we used FindAllMarkers to compare cluster against all other clusters, and FindMarkers to compare selected clusters. For each cluster, only genes that were expressed in more than 15% of cells with at least 0.15-fold difference were considered. Heatmap representations were generated as described earlier with Phantasus software (https://artyomovlab.wustl.edu/phantasus/) ^77^, using the mean expression of markers inside each cluster for each sample was used.

### Circos Plots

Circos plots depicting possible cell interactions were created using SingleCellSignalR ^81^.

### Immunohistochemistry and Confocal Imaging

To validate the findings of BAL single cell sequencing, multilabel immuno-histochemistry was performed on Naïve and SARS CoV-2 infected Rhesus macaque lungs at Day 3 and Day 14 post infection as described ^4^. The lung sections were stained for macrophages with anti-CD68 antibody, SIGLEC1 with anti-CD169 antibody, Mac_IFN_1 signature markers with anti-MX1, MX2 and ISG15 antibodies; Mac-TREM2 with anti C1q-FITC conjugated antibody and pDCs with anti-HLA-DR and anti-CD123 antibodies to validate the *in-vivo* expression of these markers in SARS CoV-2 infected lung tissue (Table S2). SARS CoV-2 nucleocapsid antibody was used to detect SARS CoV-2 and ACE-2 expression was confirmed using human anti-ACE2 antibody. DAPI was used for nuclear staining. Images were captured using Ziess LSM-800 confocal microscope.

## Statistical Analysis

Graphs were prepared and statistical comparisons were applied using GraphPad Prism v8.4.3. Statistical comparisons were performed as outlined in respective methods. One-way repeated measure ANOVA with Geisser Greenhouse correction for sphericity and Tukey post hoc correction for multiple testing (GraphPad Prism 8.4.3) was applied for statistical comparison of population clusters across timepoints as described in the figure legends. For correlation analysis, Spearman’s rank tests were applied. Statistical differences between groups were reported to be significant when the P value was less than or equal to 0.05. Data are presented as mean + standard error of mean (SEM).

## Reporting Summary

Further information on research design is available in the Nature Research Reporting Summary linked to this article.

## Data availability

All data supporting the findings of this study are available within this manuscript and its Supplementary Information. Any additional data can be requested from the corresponding authors upon reasonable request.

## Acknowledgements

The work described in this manuscript was supported by Washington University in St. Louis (S.A.K) for COVID-19 research, as well as and NIH award # R01AI123780 to S.A.K, M.M. and D.K., R01AI134236 to S.A.K. and D.K. and a COVID-19 supplement to it, an NIH award # R01AI134245 to S. M., and by institutional NIH awards P51OD111033 and U42OD010442 to the SNPRC, Texas Biomedical Research Institute. These funders had no role, however, in the design and execution of the experiments and the interpretation of data. The views expressed here are those of the authors and do not necessarily represent the views or official position of the funding agencies. The authors declare that no other financial conflict of interest exist.

