## Supplementary material for "Myeloid cell interferon responses correlate with clearance of SARS-CoV-2": Suppl Figs 1-16

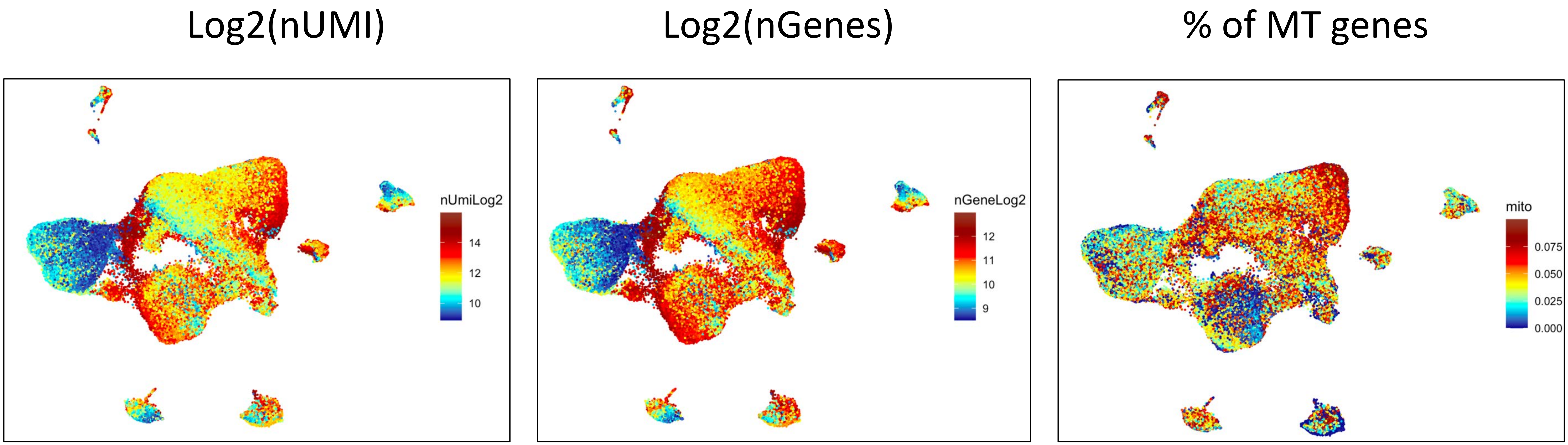
Log2(nUMI)
Log2(nGenes)
% of MT genes
nUmiLog2
14
12
10
nGeneLog2
12
11
10
9
mito
0.075
0.050
0.025
0.000

Fig S1

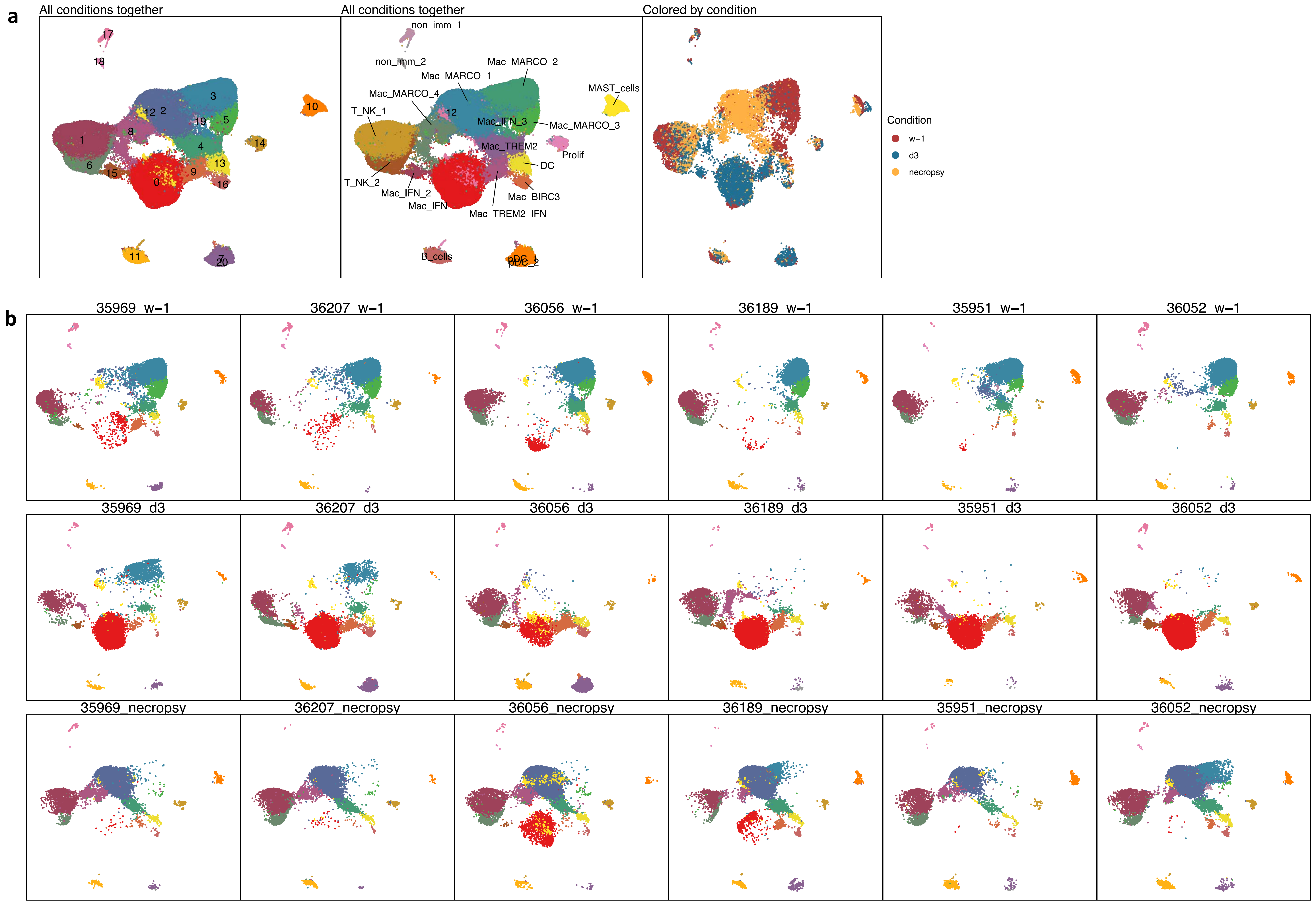
a
All conditions together
17
18
3
12
2
19
5
10
8
1
4
14
6
15
0
9
13
16
11
7
20
All conditions together
non_imm_1
non_imm_2
Mac_MARCO_2
Mac_MARCO_1
Mac_MARCO_4
MAST_cells
T_NK_1
12
Mac_IFN_3
Mac_MARCO_3
Mac_TREM2
Prolif
DC
T_NK_2
Mac_IFN_2
Mac_IFN
Mac_BIRC3
Mac_TREM2_IFN
B_cells
Colored by condition
Condition
w−1
d3
necropsy
b
35969_w−1
36207_w−1
36056_w−1
36189_w−1
35951_w−1
36052_w−1
35969_d3
36207_d3
36056_d3
36189_d3
35951_d3
36052_d3
35969_necropsy
36207_necropsy
36056_necropsy
36189_necropsy
35951_necropsy
36052_necropsy

Fig S2

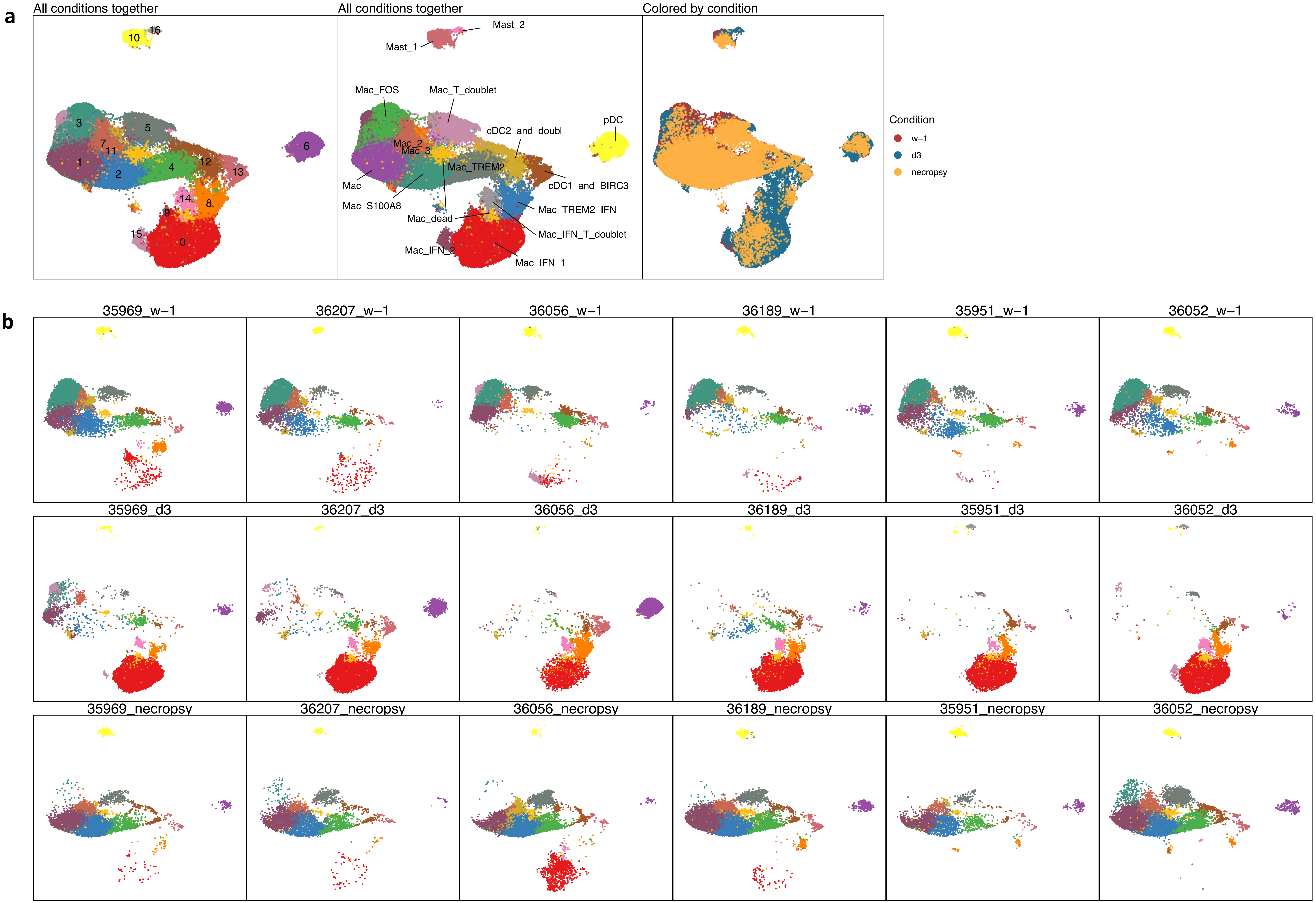
a
All conditions together
All conditions together
Colored by condition
Mast_2
Mast_1
Mac_FOS
Mac_T_doublet
cDC2_and_doubl
pDC
Mac_2
Mac_3
Mac_TREM2
cDC1_and_BIRC3
Mac
Mac_S100A8
Mac_TREM2_IFN
Mac_dead
Mac_IFN_T_doublet
Mac_IFN_2
Mac_IFN_1
Condition
w−1
d3
necropsy
b
35969_w−1
36207_w−1
36056_w−1
36189_w−1
35951_w−1
36052_w−1
35969_d3
36207_d3
36056_d3
36189_d3
35951_d3
36052_d3
35969_necropsy
36207_necropsy
36056_necropsy
36189_necropsy
35951_necropsy
36052_necropsy

Fig S3

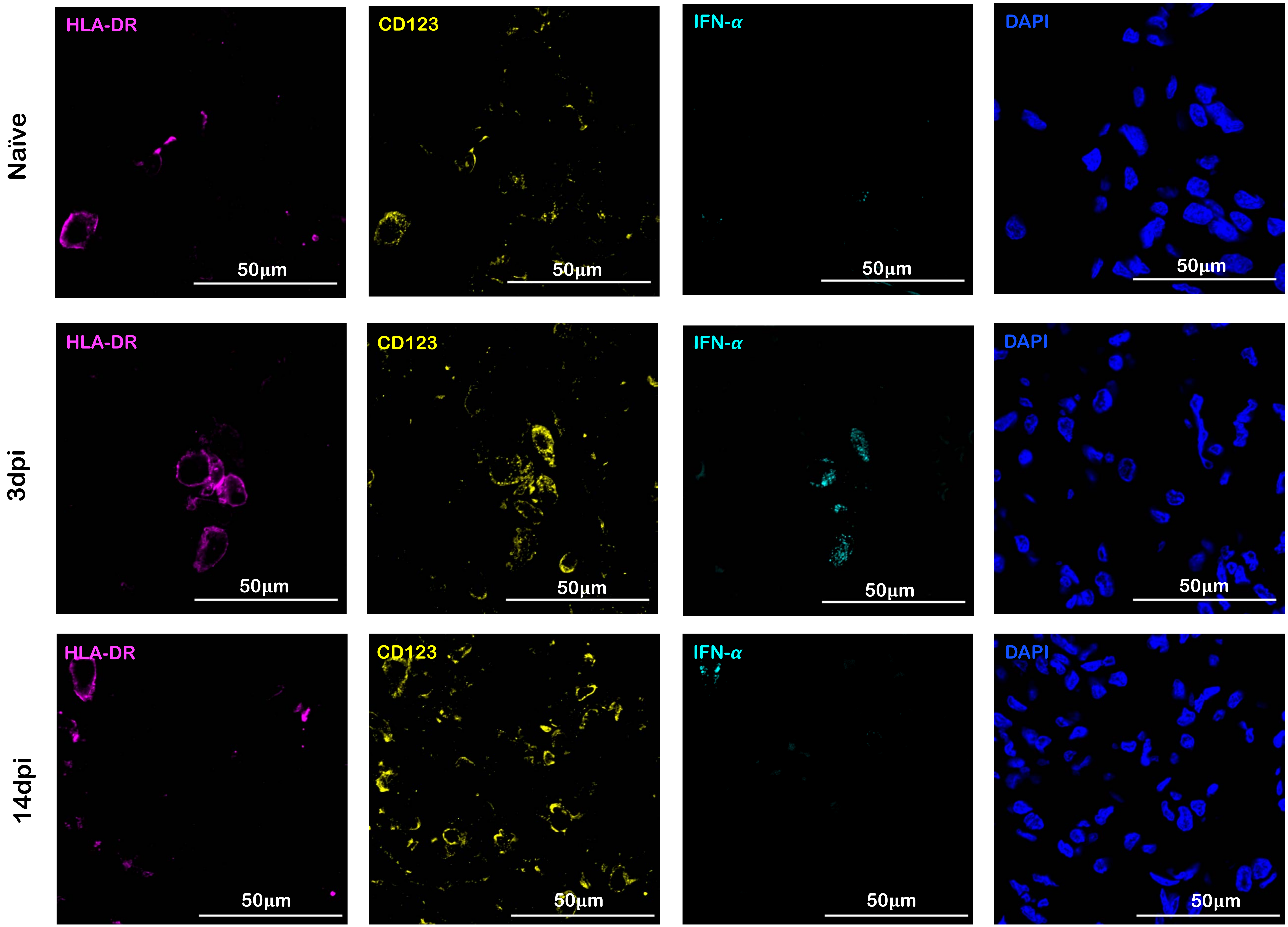


Fig S4

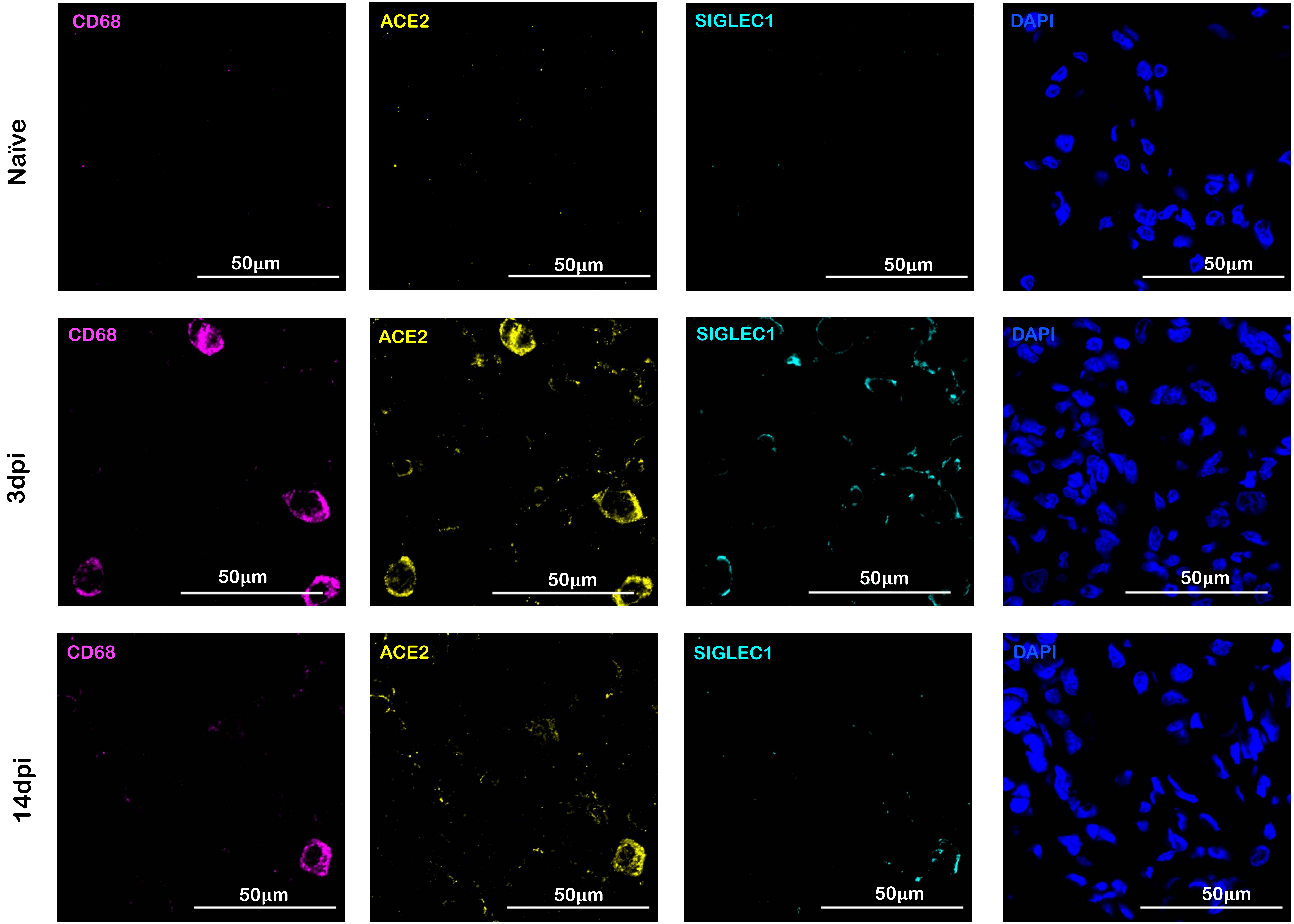


Fig S5

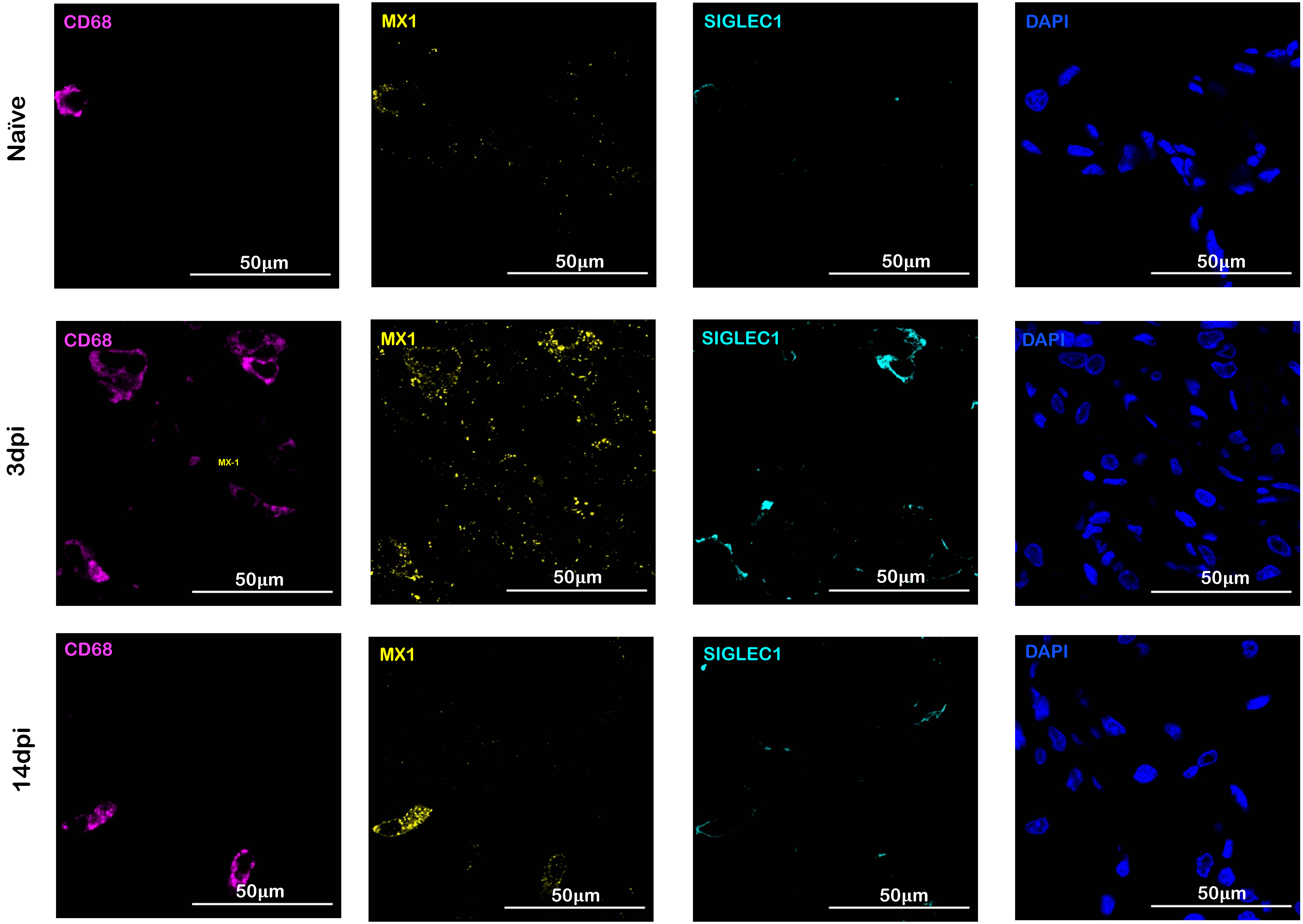


Fig S6

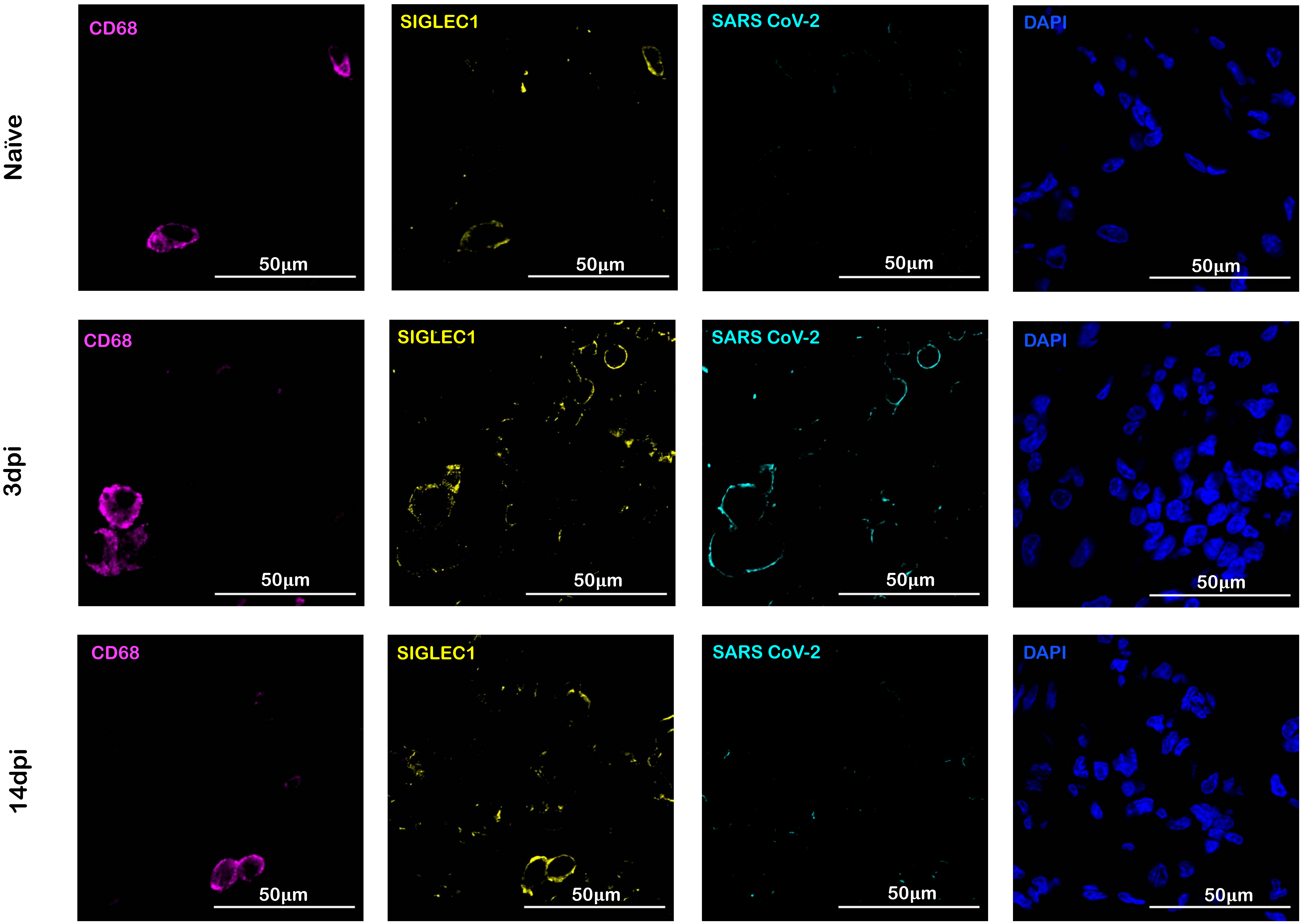


Fig S7

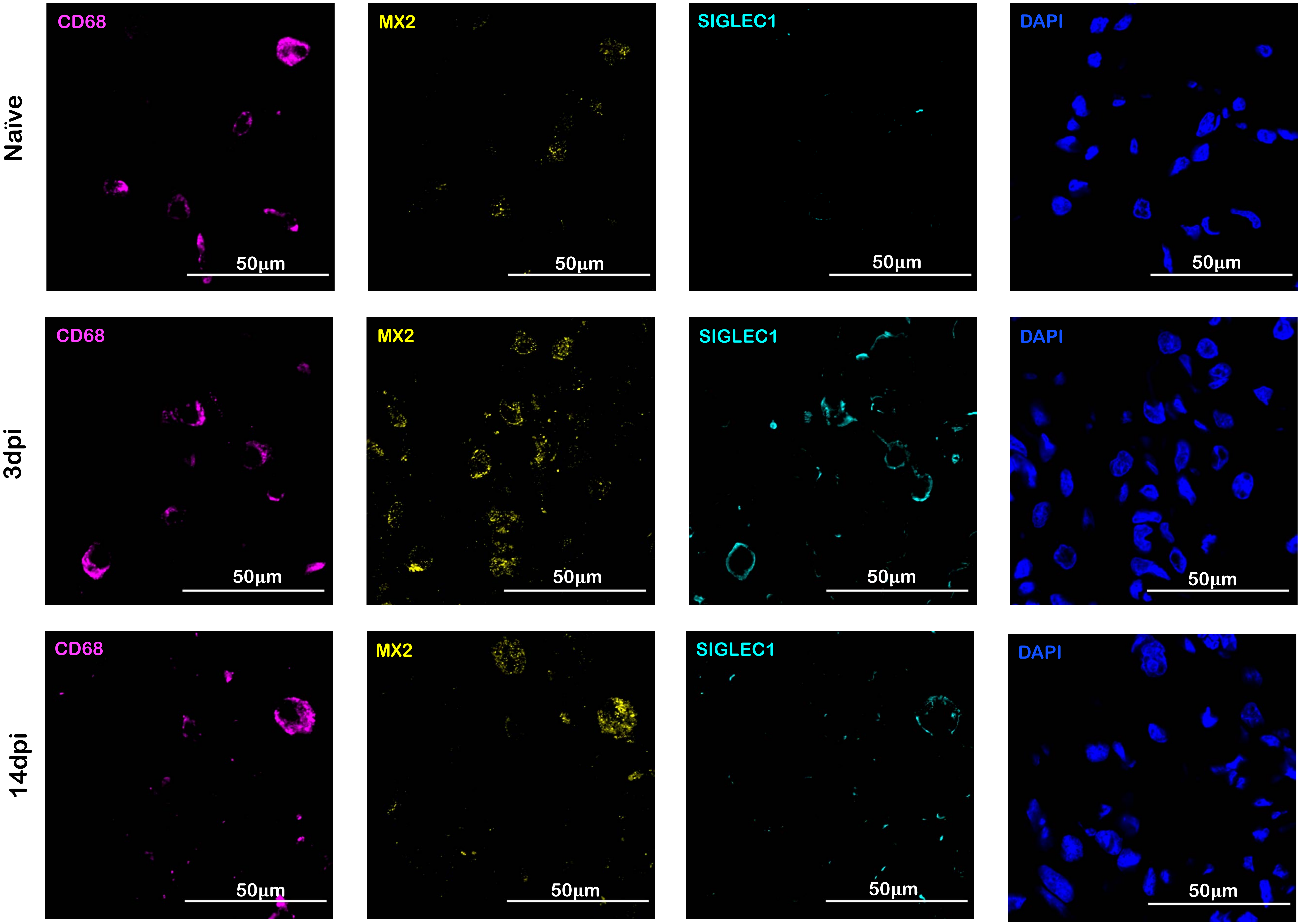


Fig S8

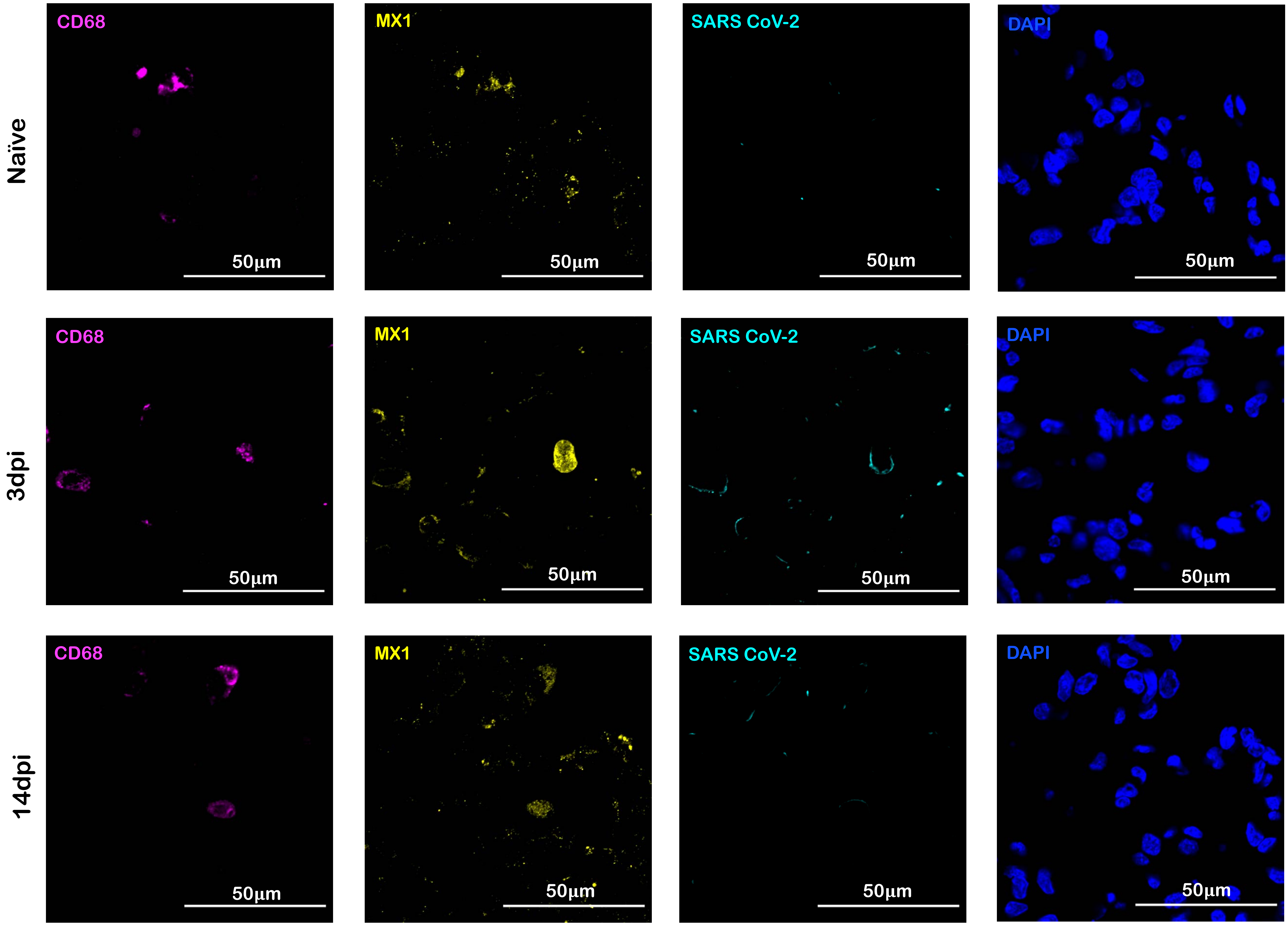


Fig S9

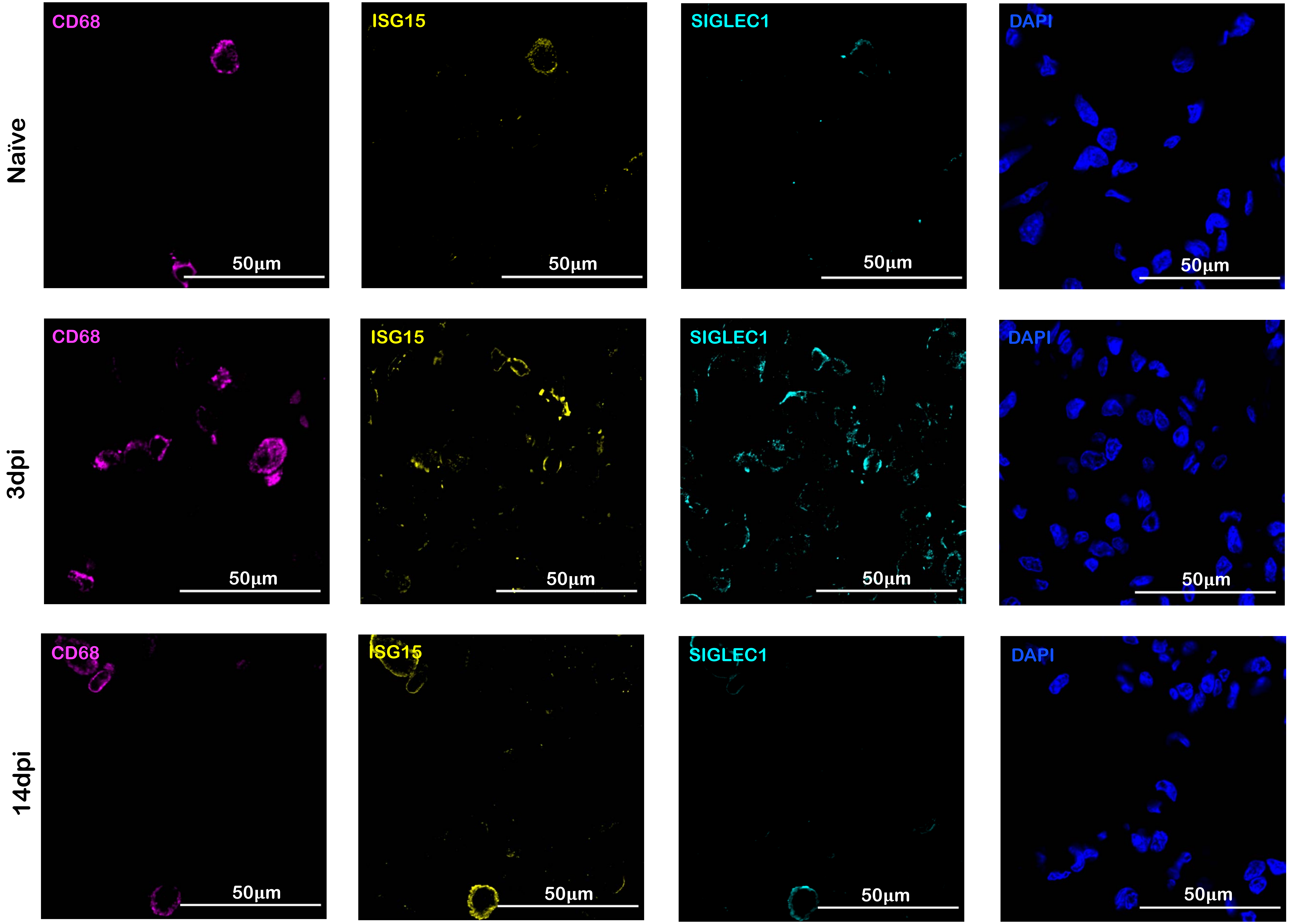


Fig S10

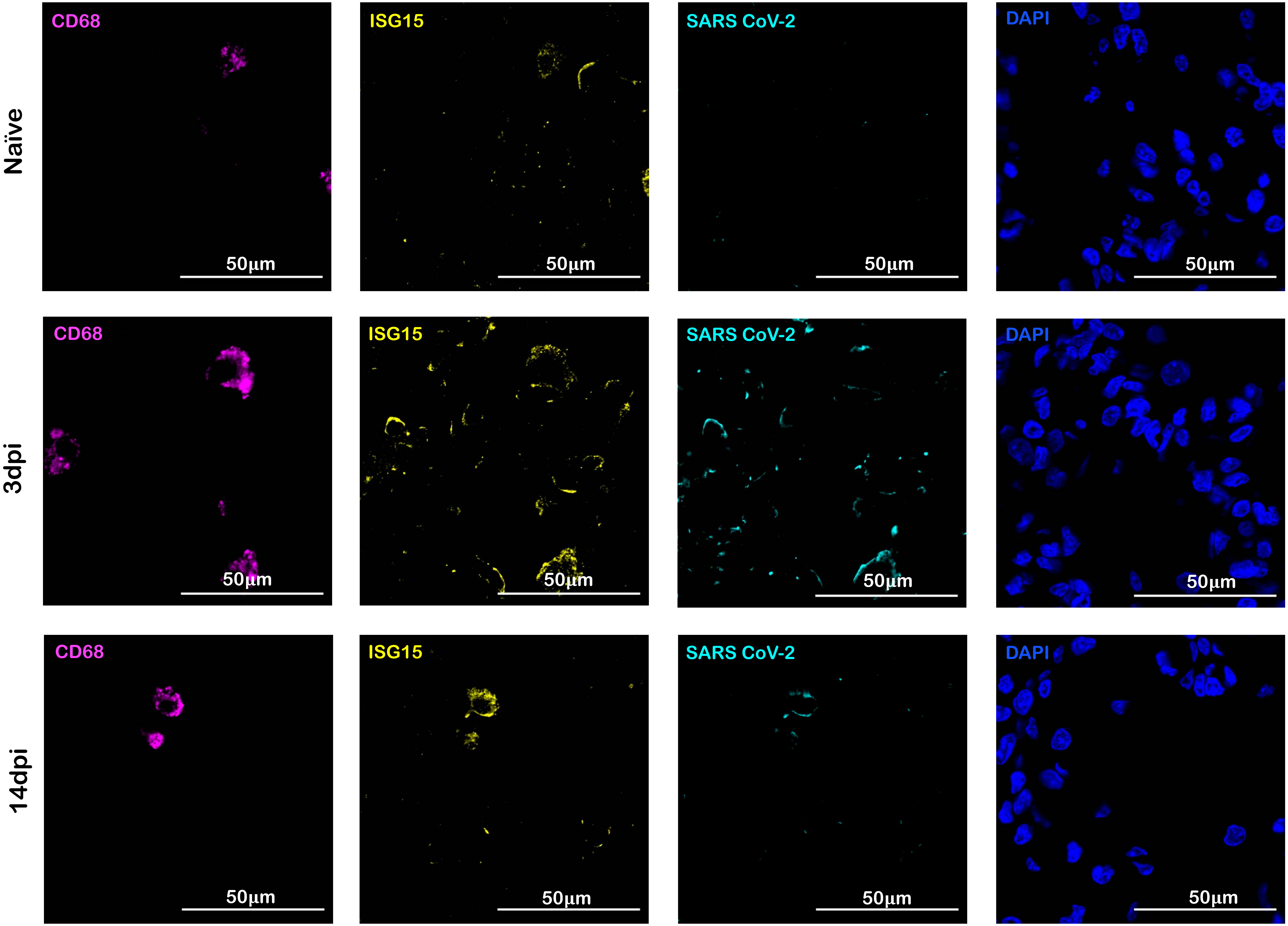


Fig S11

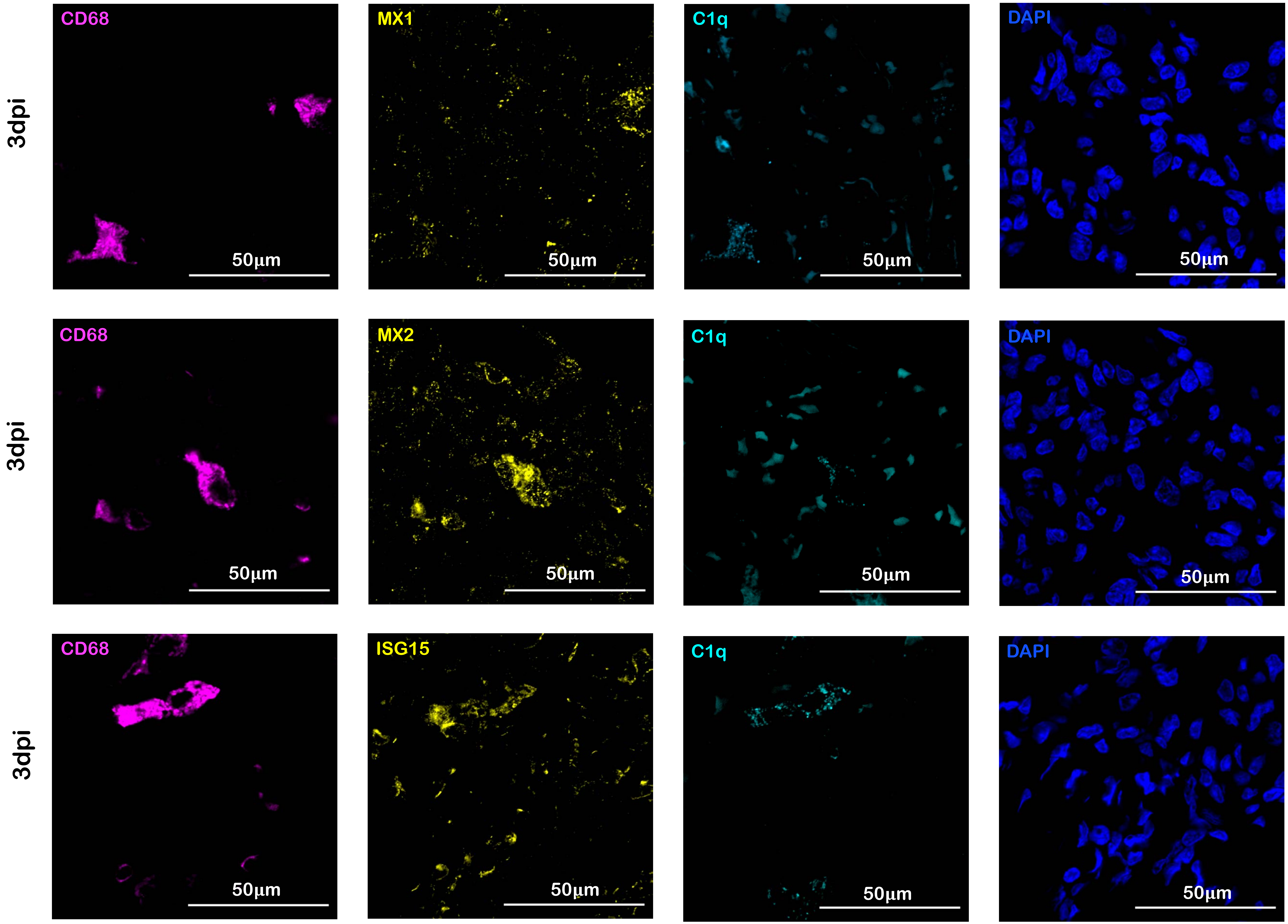
3dpi
CD68
50µm
MX1
50µm
C1q
50µm
DAPI
50µm
3dpi
CD68
50µm
MX2
50µm
C1q
50µm
DAPI
50µm
3dpi
CD68
50µm
ISG15
50µm
C1q
50µm
DAPI
50µm

Fig S12

**a**

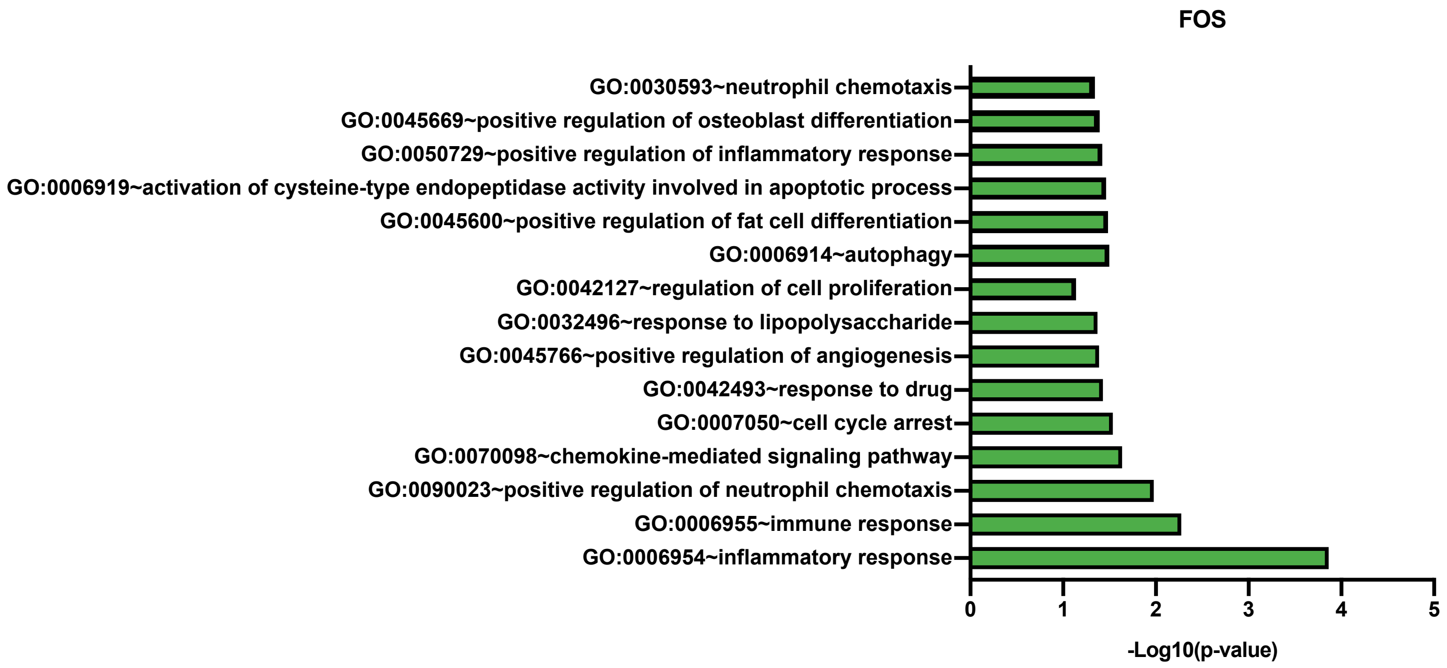
FOS
GO:0030593~neutrophil chemotaxis
GO:0045669~positive regulation of osteoblast differentiation
GO:0050729~positive regulation of inflammatory response
GO:0006919~activation of cysteine-type endopeptidase activity involved in apoptotic process
GO:0045600~positive regulation of fat cell differentiation
GO:0006914~autophagy
GO:0042127~regulation of cell proliferation
GO:0032496~response to lipopolysaccharide
GO:0045766~positive regulation of angiogenesis
GO:0042493~response to drug
GO:0007050~cell cycle arrest
GO:0070098~chemokine-mediated signaling pathway
GO:0090023~positive regulation of neutrophil chemotaxis
GO:0006955~immune response
GO:0006954~inflammatory response
0
1
2
3
4
5
-Log10(p-value)

**b**

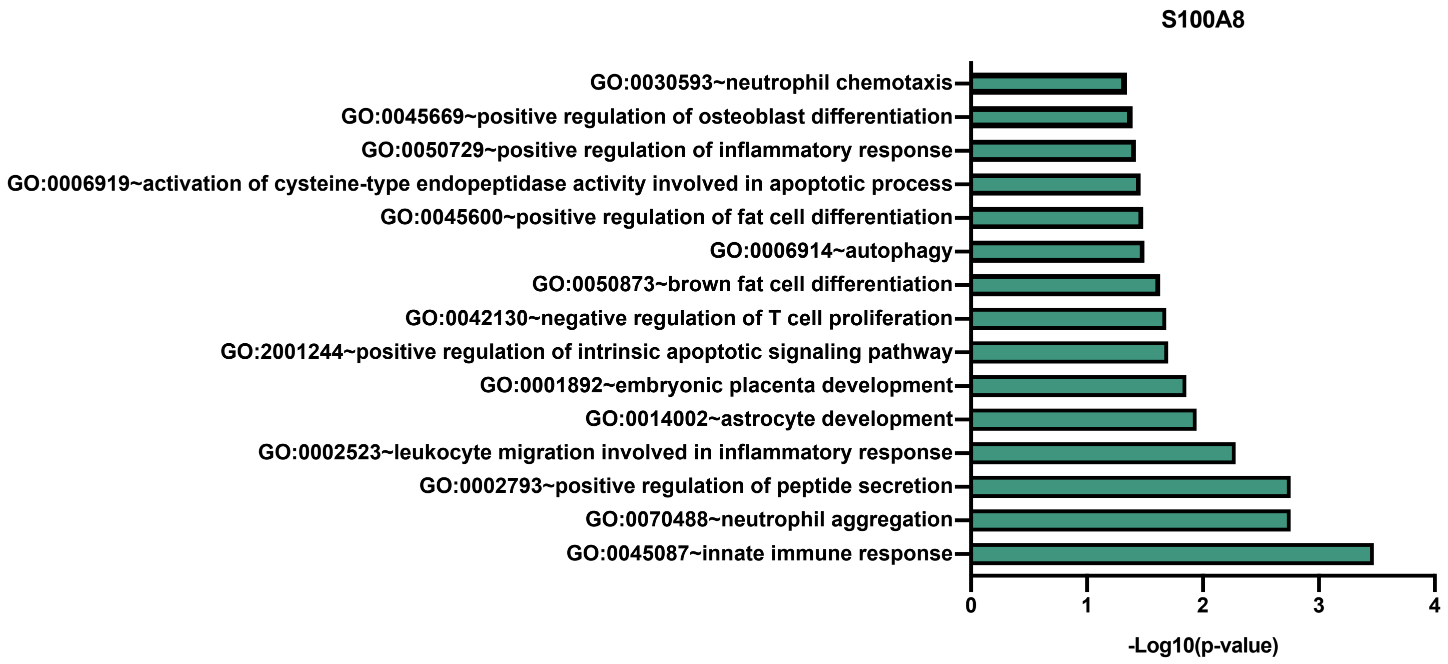
S100A8
GO:0030593~neutrophil chemotaxis
GO:0045669~positive regulation of osteoblast differentiation
GO:0050729~positive regulation of inflammatory response
GO:0006919~activation of cysteine-type endopeptidase activity involved in apoptotic process
GO:0045600~positive regulation of fat cell differentiation
GO:0006914~autophagy
GO:0050873~brown fat cell differentiation
GO:0042130~negative regulation of T cell proliferation
GO:2001244~positive regulation of intrinsic apoptotic signaling pathway
GO:0001892~embryonic placenta development
GO:0014002~astrocyte development
GO:0002523~leukocyte migration involved in inflammatory response
GO:0002793~positive regulation of peptide secretion
GO:0070488~neutrophil aggregation
GO:0045087~innate immune response
0
1
2
3
4
-Log10(p-value)

***Fig S13***

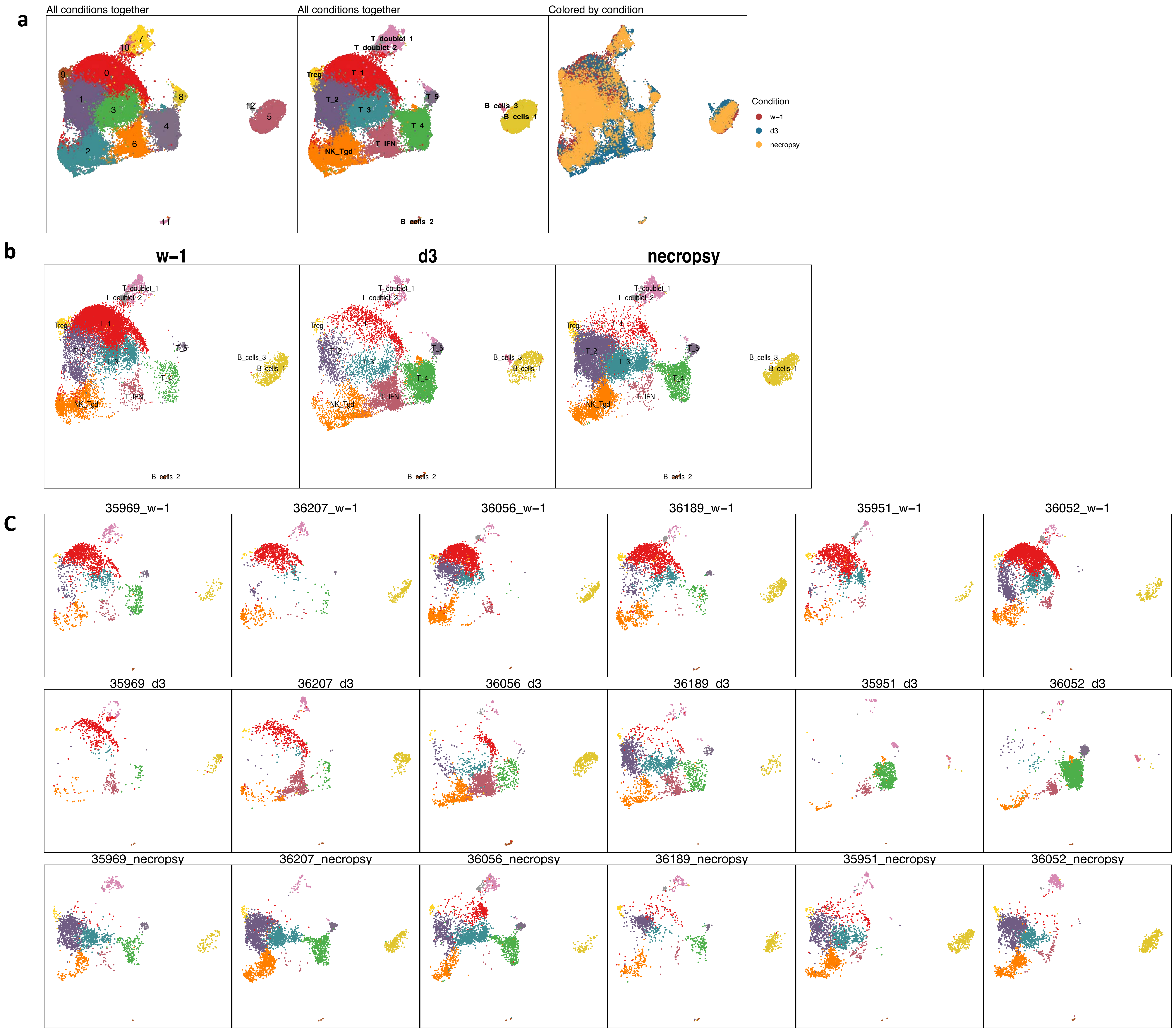
a
All conditions together
All conditions together
Colored by condition
Condition
w−1
d3
necropsy
T_doublet_1
T_doublet_2
Treg
T_1
T_2
T_3
T_4
T_5
T_IFN
NK_Tgd
B_cells_1
B_cells_2
B_cells_3
b
w−1
d3
necropsy
c
35969_w−1
36207_w−1
36056_w−1
36189_w−1
35951_w−1
36052_w−1
35969_d3
36207_d3
36056_d3
36189_d3
35951_d3
36052_d3
35969_necropsy
36207_necropsy
36056_necropsy
36189_necropsy
35951_necropsy
36052_necropsy

Fig S14

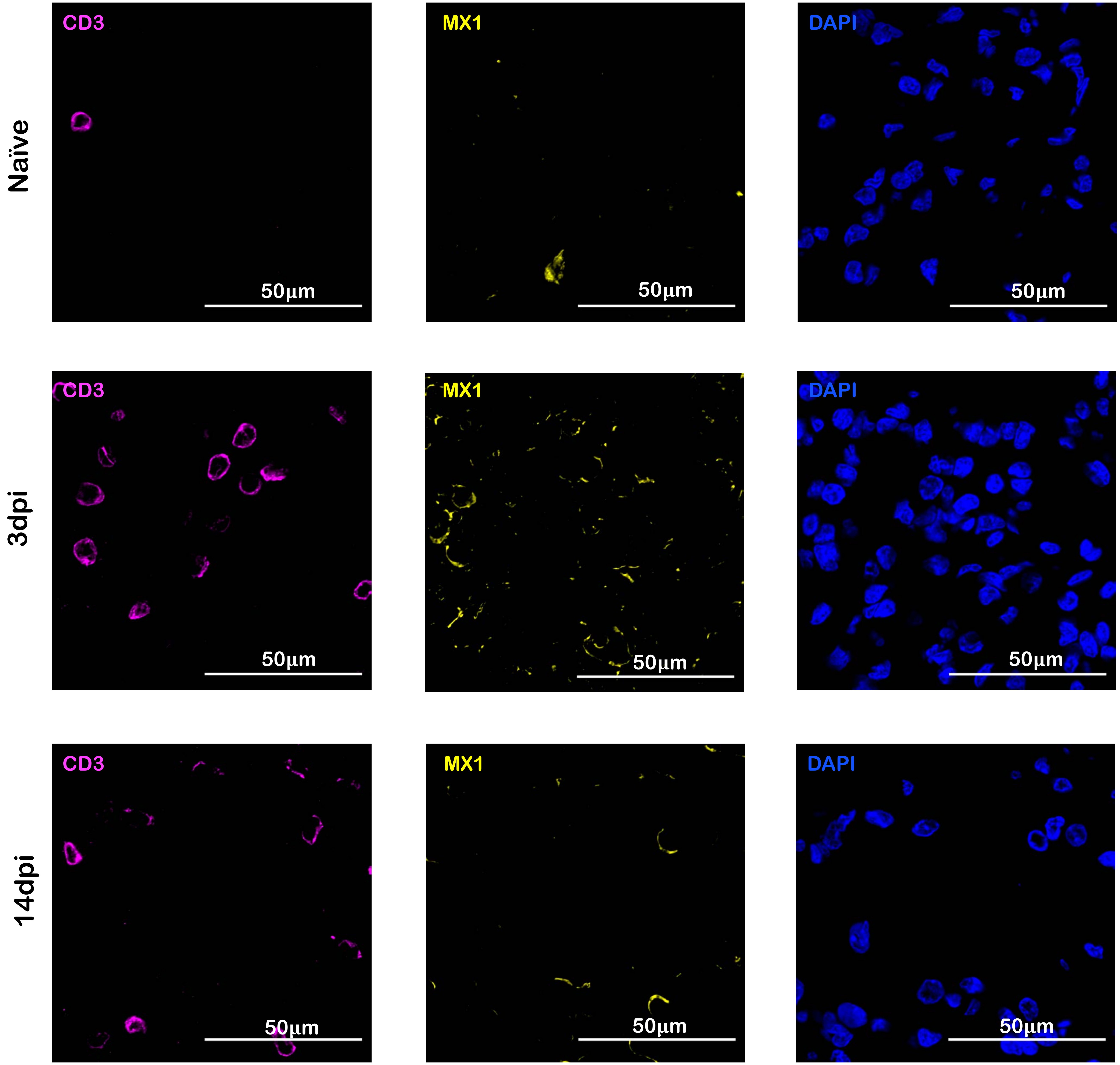


Fig S15

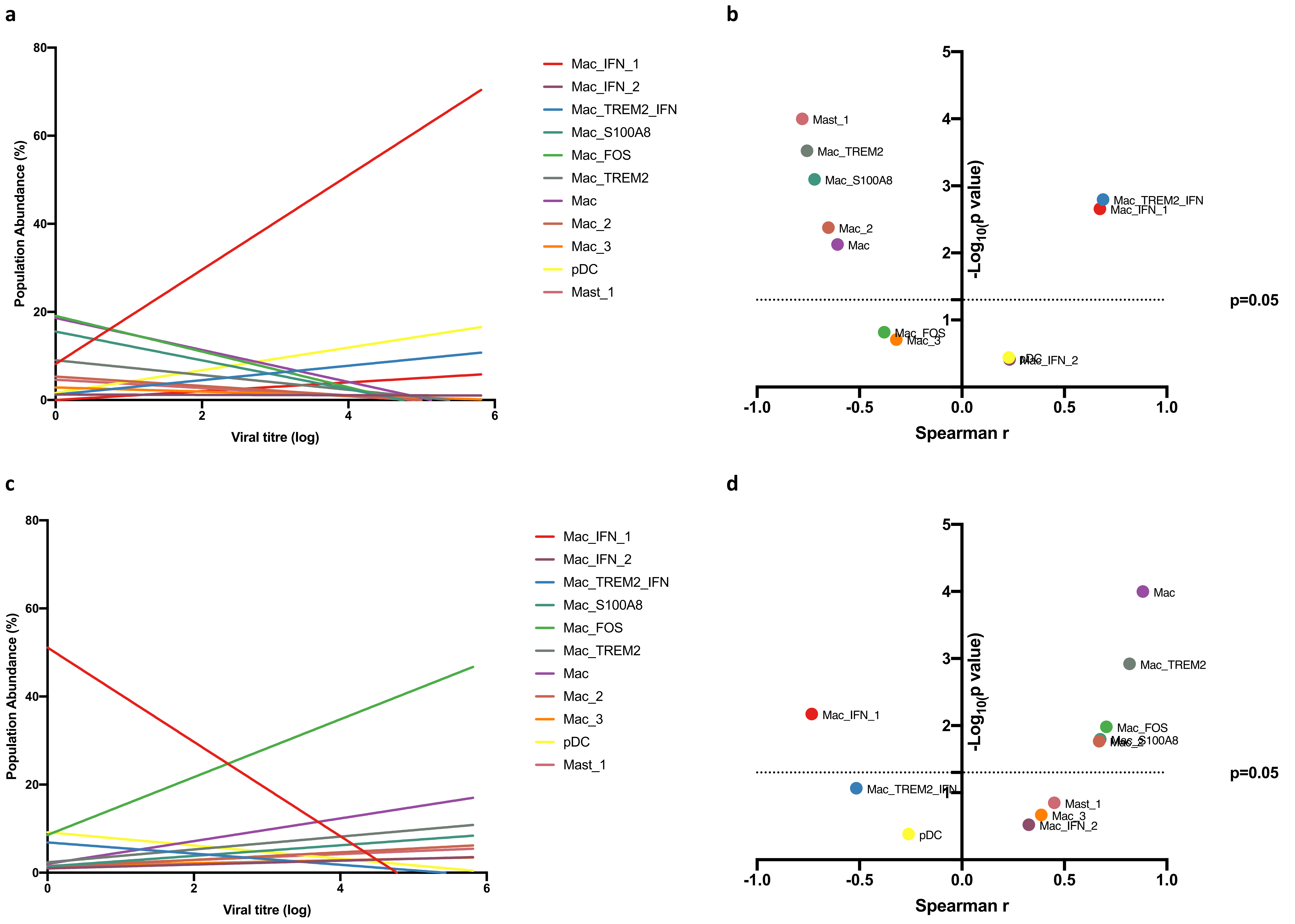
a
b
c
d
Population Abundance (%)
Viral titre (log)
Mac_IFN_1
Mac_IFN_2
Mac_TREM2_IFN
Mac_S100A8
Mac_FOS
Mac_TREM2
Mac
Mac_2
Mac_3
pDC
Mast_1
-Log10(p value)
Spearman r
p=0.05

Fig S16
